# Effects of frugivore species pool and seed size on the diversity and functional composition of frugivores visiting fruiting trees

**DOI:** 10.1101/2025.01.31.635888

**Authors:** Rohit Naniwadekar, Abhishek Gopal, Rintu Mandal, Arpitha Jayanth, Vatcharavee Sriprasertsil, Sartaj Ghuman, Navendu Page, Saniya Chaplod, Himanshu Lad, Aditya Gadkari, Vignesh Chandran, Natasha Desai, Rasika Kadam, Bee Choo Strange, Vijak Chimchome, George A. Gale, Jahnavi Joshi

## Abstract

The relationship between biodiversity and ecosystem functioning in seed dispersal remains understudied despite its critical role in maintaining plant diversity in the tropics. Field studies on this relationship are often confounded by environmental and phylogenetic variations across species richness gradients. We examined how overall avian frugivore species richness at a site influenced the frugivore richness, visitation rates and functional composition of two key effect traits⸺beak width and hand-wing index⸺on fruiting trees. Across six sites in tropical Asia, spanning a sevenfold gradient in frugivore species richness but with similar forest types and phylogenetically nested frugivore communities, we recorded 34,014 interactions between 134 avian frugivores and 131 plant species. Our results provide some support for the biodiversity-ecosystem functioning relationship, as higher overall frugivore species richness increased the number of frugivore species visiting individual fruiting trees but not the functional composition of frugivores. Seed size had a stronger influence on the frugivore species richness, visitation rates, and the beak size of visiting frugivores, highlighting the dominant role of morphological trait matching in influencing plant-frugivore interactions. Our findings suggest functional redundancy in certain aspects of seed dispersal effectiveness due to density compensation and the presence of key seed disperser lineages in species-poor sites.

## 1. BACKGROUND

Wet tropical forests are characterised by intricate interactions between plants and frugivores, which are critical in maintaining plant diversity. Understanding how frugivore diversity impacts seed dispersal outcomes for plants has intrigued biologists given its implications for ecosystem functioning and resilience, especially due to global change. While the positive impact of frugivore abundance on seed dispersal is well documented [1], the influence of frugivore diversity on plants’ seed dispersal outcomes remains mixed. For example, Garcia and Martinez [2] found that thrush species richness positively affected seed dispersal, whereas Quitian et al., [3] found that the functional diversity of frugivores did not directly influence the functional diversity of removed fruits, a potential outcome of functional complementary [4]. Despite three decades of research on biodiversity-ecosystem functioning relationships [5–7], our understanding of the relationship between frugivore diversity and seed dispersal remains limited [8]. This knowledge is crucial for understanding functional redundancy and complementarity among seed dispersers, especially as islands and defaunated sites tend to have lower diversity of frugivores [9,10].

The interactions of frugivores and fleshy-fruited plants are mediated by seed traits [9,11], wherein smaller seeded plants are more likely to have functional redundancy due to greater diversity of frugivores visiting them compared to larger seeded plants that might be more specialised with respect to the frugivore community visiting them. Seed traits significantly mediate the relationship between frugivores and plants through morphological trait matching. Small-seeded plants, which typically attract more frugivore species, exhibit higher functional redundancy compared to large-seeded plants [12]. This underscores the importance of factoring seed traits while studying the effects of frugivore diversity on seed dispersal.

Existing studies on the impact of frugivore diversity on seed dispersal are predominantly conducted along disturbance gradients [2], focused on a few plant species [12,13], or situated in species-poor temperate habitats [2,14]. There is a notable lack of information from hyperdiverse wet tropical forests, which harbour high frugivore diversity but face significant threats. Moreover, since existing studies focus on a few plant species [13], community-wide perspectives remain relatively understudied.

Field-based studies examining the relationship between biodiversity and ecosystem functioning are often influenced by multiple drivers, including environment, species composition and diversity, with covarying drivers often influencing each other [7,15]. One of the main criticisms of such studies is that environmental conditions (e.g., habitat type), which influence ecosystem functioning, are often not controlled. Studies conducted across broad geographical scales frequently encounter challenges due to varying environmental conditions and phylogenetic relationships of frugivores. Moreover, traits are influenced by environment and phylogenetic relationships, posing issues for discerning the main effects due to confounding influences of varying environments and the evolutionary histories of communities [16]. Studies across ‘species richness anomalies’ that control for environmental gradients and phylogenetic relationships of species are limited [16].

In wet tropical forests, frugivorous birds play a pivotal role in dispersing seeds of diverse plant species [17]. These ecosystems host high bird diversity, and are characterised by significant variations in bird functional traits [18]. Among avian frugivores, two traits are particularly relevant for seed dispersal: beak width, which determines the size of fruits and seeds that a bird can consume and disperse without causing seed damage, and hand-wing index, which reflects the bird’s dispersal ability [3,19,20]. Birds with greater dispersal abilities can locate patchily distributed fruit resources and disperse seeds away from the parent plant, occasionally facilitating long-distance seed dispersal, which is critical in the context of habitat fragmentation and climate change [21]. While functional diversity is more strongly associated with ecosystem functioning than taxonomic diversity [7], we were more interested in the functional composition of frugivores visiting fruiting trees, as these traits are known to affect seed dispersal outcomes.

Moreover, we aimed to determine how predictors influence the species richness of birds visiting individual fruiting trees and their visitation rates. The richness of birds visiting fruiting tree can affect the qualitative component of seed dispersal, as different species may contribute differentially due to behavioural differences [22]. Visitation rates represent a key quantitative component of seed dispersal, as the number of frugivores is often directly linked to the number of seeds dispersed. Understanding how the frugivore species pool influences these patterns can provide insights into the prevalence of density compensation in species-poor sites, an aspect that is relatively understudied.

Given this background, we conducted a field study across six wet tropical forest sites in the Oriental realm, spanning a sevenfold gradient in avian frugivore species richness ranging from seven to 48 frugivore species. The diversity across six sites has been shaped by distinct biogeographic processes that allowed us to examine on the natural gradient in species richness. The environmental conditions, like annual precipitation and forest types (tropical evergreen forests), are similar across these sites (Fig. 1). All sites had representation from most of the main lineages of seed dispersers in the region viz-a-viz, Bucerotids (hornbills), Pycnonotids (bulbuls), Megalaimids (barbets), Columbids (Pigeons) and Sturnids (mynas) [19,23,24] (Fig. 1). We documented interactions between fruit-eating birds and fleshy-fruited plants and examined the relative effects of overall frugivore species richness and seed size on the taxonomic and functional composition of frugivores visiting individual fruiting trees. Following the expectations of biodiversity-ecosystem functioning relationships, we hypothesised there would be a positive relationship between the taxonomic diversity of frugivores visiting fruiting trees with overall frugivore species richness. Given density compensation [25], we did not expect any relationship between the visitation rates of frugivores on fruiting trees with overall frugivore species richness. Given that complementary effects would facilitate the packing of greater diversity in the community, we hypothesised that seed size would have a stronger influence than overall frugivore species richness on taxonomic diversity and the community-weighted mean of beak width due to morphological trait matching [11]. While body size does not always correlate with dispersal ability [26], large-seeded plants are primarily dispersed by large-bodied frugivores, which are likely to have greater dispersal ability [27,28]. Thus, we also expected dispersal ability to be closely associated with seed traits.

**Figure 1.**
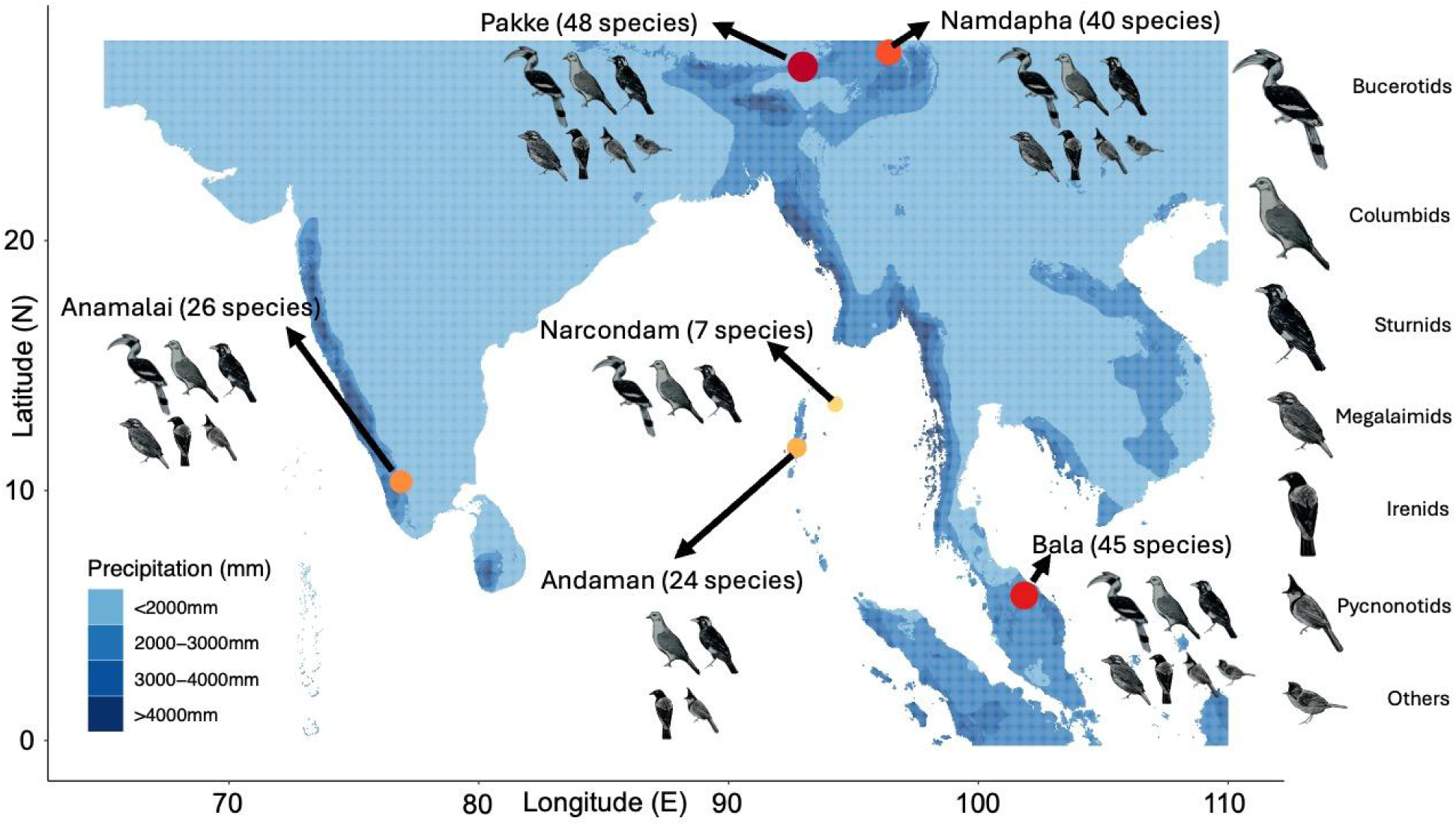
Map showing six sites across four biodiversity hotspots in South and South-east Asia, differing in frugivore species richness. All sites had similar habitats (evergreen forests with > 2000 mm annual rainfall) and nested frugivore assemblages. Site names are shown along with the frugivore species pool recorded at each site in parentheses. In the figure, ‘others’ refers to bird families like Leiothrichidae or Calyptomenidae, which were exclusively represented in the species-rich sites.

## 2. METHODS

The study was conducted across six sites spread across four biodiversity hotspots - Western Ghats-Sri Lanka, Eastern Himalaya, Indo-Myanmar and Sundaland in the Oriental realm. All the sites are tropical lowland evergreen rainforest sites below 1,200 m above sea level and receive more than 2000 mm of annual rainfall (Fig. 1) [29]. These sites within the Oriental realm represent distinct biogeographic subunits, suggesting distinct biogeographic processes may have shaped the diversity of frugivore assemblages. The frugivore communities on the volcanic Narcondam Island, the continental South Andaman Island, and the Western Ghats is most likely shaped by dispersal. On the other hand, Pakke and Namdapha in Eastern Himalaya and Bala in peninsular Thailand could be a product of in-situ speciation, dispersal and vicariance. The most species-poor site was the small, volcanic, oceanic island of Narcondam, which had seven avian frugivore species, while the species-rich sites in the Eastern Himalaya, Indo-Myanmar and Sundaland regions had more than 40 species of frugivores (Table S1). Thus, biogeography allowed us to have these sites with similar tropical evergreen forest types and nested phylogenetic composition of frugivores thereby minimising the influence of environmental factors and phylogenetic variations on observed patterns.

We observed 544 individual trees of 131 plant species across the six sites (Table S2). The tree watches were conducted following established protocols with two or three observers watching fruiting trees and recording species identity and the number of frugivorous birds that arrived on fruiting trees in early mornings and/or late afternoons coinciding with activity patterns of the frugivores [19]. The total tree watch sampling effort was 4,246.7 hr. Across five of the six sites, the duration of the tree watch was kept fixed to six hours in the first half of the day. In one of the first sites, we sampled an individual tree for a full day. The mean duration of the tree watches was 7.8 hr (SD = 2.5 hr; median = 6 hr; range: 4.2–13.3 hr). However, the duration of the tree watch did not influence the richness of frugivores visiting fruiting trees (Fig. S1). The number of tree species observed at each site ranged from 13 to 48, but it was not correlated with the number of frugivore species observed (*ρ* = 0.64; *S* = 12.679; *p* = 0.1731) (Table S2).

Since we did not have precise seed width information for some species, we classified the seeds into small (< 5 mm), medium (5–15 mm) and large (> 15 mm) following existing literature [19] (Table S3). We had representation of small-, medium- and large-seeded plants across all the sites (Fig. S2).

### Statistical Analysis

All analyses were done in R ver. 4.3.3 [30].

To determine whether the composition of avian frugivores in species-poor communities was a nested subset of the species-rich communities, we generated a matrix of six sites and 26 bird families. We used the ‘oecosimu’ function of the R package ‘vegan’ to estimate NODF (Nestedness Overlap and Decreasing Fill) for the observed community. We generated 1000 null communities using the method ‘r1’, which ensures that the row and column totals are preserved. We compared the observed metric to the null to determine whether the observed community was significantly more nested than the null communities.

We used the function ‘estimateD’ of R package ‘iNEXT’ to determine sampling coverage based on abundance data from individual-focal trees and compare the overall richness of frugivores visiting fruiting trees in different seed size classes at each site. This analysis was done for each site separately. Sampling coverage measures how well the community has been sampled and is among the fairest measures to standardise and compare diversity across communities [31].

We used the function ‘glmmTMB’ as implemented by package ‘glmmTMB’ to fit a generalised linear mixed model (negative binomial error structure since the Poisson model was overdispersed) for determining the influence of seed size and overall avian frugivore richness in the community on the number of avian frugivore species visiting a fruiting tree [32]. We centered the overall species richness to reduce the correlation between the coefficients of fixed effects in the model. We used the site as a random effect to measure unmeasured heterogeneity across sites. We determined marginal-*R^2^_marginal_* to estimate the percentage variation explained by the model. To determine the relative influence of seed size and overall frugivore species richness in explaining the variation in taxonomic diversity, we separately estimate marginal *R^2^_marginal_* with a seed size-only model and overall species richness-only model.

We used the ‘lmer’ function as implemented by the package ‘lmertest’ to fit a linear mixed model (Gaussian error structure) and determine the influence of seed size and overall avian frugivore richness on the visitation rates of frugivores and the community-weighted mean of beak width and hand-wing index [33]. We used the natural logarithms of visitation rate (per hour) of avian frugivores on individual trees to estimate the community-weighted mean. We used the site as a random effect. We used the ‘functcomp’ function as implemented by package ‘FD’ in R to estimate the community-weighted mean[34].

## 3. RESULTS

We documented 34,014 interactions between 138 avian frugivores and 131 species of plants across the six sites. The sampling coverage of small, medium, and large-seeded plants for each of the six sites was > 0.99, except for large-seeded plants in Bala, which had a coverage of 0.94, indicating sampling adequacy (Table S4). Interestingly, despite being on the mainland, the number of frugivore species in Anamalai (southern Western Ghats; 26 species) was similar to the continental island site (south Andamans; 24 species) and was almost half of the most species-rich site, Pakke in the Eastern Himalaya (48 species). The observed community was significantly more nested than expected by null (NODF_obs_ = 67.1, NODF_null_ = 58.6 (95% CI: 48.9 – 66.5), *p* = 0.035; Standardised Effect Size = 1.91).

### Taxonomic diversity across sites and seed size classes

We found that the number of avian frugivore species that visited the fruiting tree differed across the three seed size classes and was positively associated with the overall number of avian frugivore species in the community (Fig. 2A and B; Table S5). The variation explained by the fixed effects (*R^2^_marginal_*) was 48.2%. Variation explained by the fixed effects in the model with only seed size as the predictor was 37%. In comparison, the *R^2^_marginal_* was 14.9% in the model with only overall species richness as the predictor, highlighting the greater influence of seed traits on frugivore richness on fruiting trees across the sites than the overall avian frugivore richness in the community.

**Figure 2.**
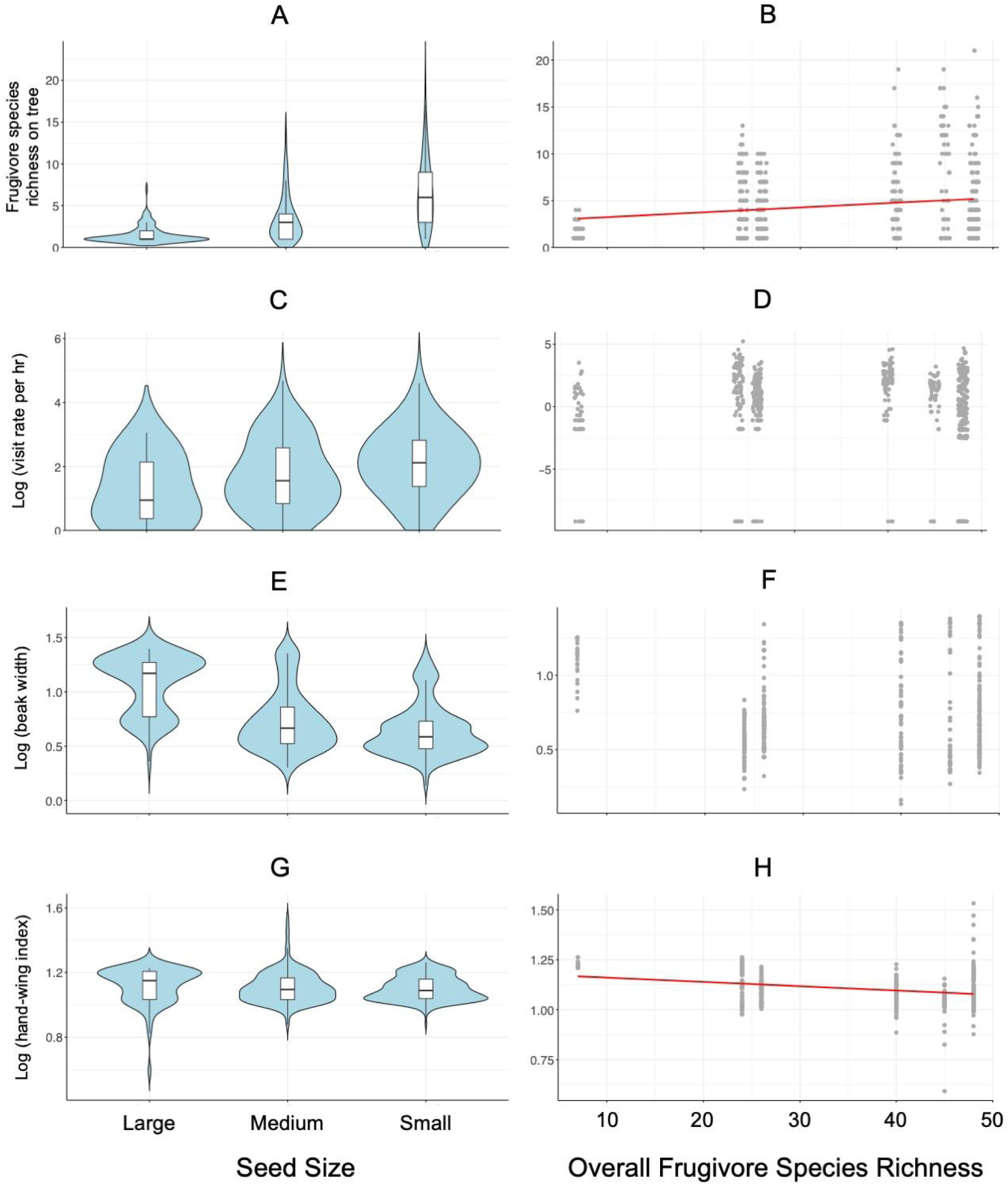
The violin plots with nested box plots show the relationship between seed size classes (small: < 5 mm; medium: 5-15 mm; large: > 15 mm) and the number of frugivore species visiting a fruiting tree (A), visitation rate of frugivores per hour (C), and log of community-weighted means of beak width (E) and hand-wing index (G). The scatter plots show the relationship between overall frugivore species richness at the site and the number of frugivore species visiting a fruiting tree (B), visitation rate of frugivores per hour (D), and log of community-weighted means of beak width (F) and hand-wing index (H). The points have been jittered to minimise overplotting and improve visibility.

### Frugivore visitation rates across sites and seed size classes

We found that the visitation rates of frugivorous birds were significantly higher for medium- and small-seeded plants than for large-seeded plants (Fig. 2C; Table S6). We did not find support for the relationship between the visitation rates of frugivorous birds and overall frugivore species richness in the community (Fig. 2D; Table S6). The *R^2^_marginal_* was 0.25.

### Beak width and hand-wing index across sites and seed classes

We found that the community-weighted mean of beak width of frugivorous birds was significantly smaller for medium- and small-seeded plants than for large-seeded plants (Fig. 2E; Table S7). We did not find support for the relationship between the community-weighted mean of beak width and overall frugivore species richness in the community (Fig. 2F; Table S7). The *R^2^_marginal_* was 0.24.

We found that the community-weighted mean of the hand-wing index of frugivorous birds was not associated with seed size (Fig. 2G; Table S8). However, it exhibited a statistically significant negative association with overall species richness, indicating that birds that visited fruiting trees in the species-rich sites tended to have lower dispersal ability than those in species-poor sites (Fig 2H; Table S8). The *R^2^_marginal_* was 0.20.

## 4. DISCUSSION

This study contributes to the long-standing research interest in understanding the relationship between biodiversity and ecosystem functioning by examining links between frugivore richness at a site and multiple quantitative and qualitative measures of seed dispersal. These measures include the number of frugivores visiting a fruiting tree, their visitation rates, and the abundance-weighted composition of two key effect traits associated with avian seed dispersal: the beak width and dispersal ability. To minimise confounding factors, the study controlled for environmental influences by sampling exclusively in evergreen tropical forests and addressed phylogenetic constraints by sampling phylogenetically nested frugivore communities within south and south-east Asia. We found evidence supporting the biodiversity-ecosystem functioning hypothesis, as greater diversity of frugivores in the community influenced the richness of frugivores visiting fruiting trees. However, greater frugivore richness did not necessarily enhance the functional composition of frugivores in terms of foraging or dispersal abilities. Our results provide strong evidence for morphological trait matching between seeds and frugivores as seed size exerted a stronger influence than overall frugivore richness on key metrics, including the number of frugivores visiting a fruiting tree, their visitation rates, and the beak size of birds that interacted with fruits. This finding suggests that morphological compatibility overrides the broader influence of site-level frugivore richness in shaping plant-frugivore interactions, which has implications for functional redundancy and complementarity, as discussed below.

Visitation rates, which positively influence seed dispersal effectiveness [35,36], are a robust measure of the quantitative role of seed dispersal outcomes. However, species abundance does not necessarily correlate with species richness due to density compensation [37]. Our findings reveal that visitation rates on individual trees were not influenced by the overall frugivore species richness at the site, suggesting density compensation in species-poor sites. Narcondam Island, the most species-poor site in this study, has the highest reported densities of hornbills globally [10], pointing towards density compensation in the absence of frugivore species. Additionally, Narcondam Island hosts the highest densities of *Ficus* and other hornbill food plants [10], underscoring the key role that few frugivore species can potentially play in shaping the plant communities through seed dispersal.

The relative roles of functional redundancy (or similarity) and complementarity in biodiversity-ecosystem functioning literature are debated. Our species-poor community has only seven frugivore species which is comparable to some temperate sites. However, we failed to discern patterns in beak width with increasing species richness, suggesting functional redundancy in species-rich communities. This is a likely consequence of the way these communities are assembled. Despite biogeographic influences in shaping the frugivore diversity, every community included key representatives of large-bodied frugivores (e.g., Bucerotids or Columbids) and small-bodied frugivores (e.g., Sturnids, Pycnonotids, Irenids, or Megalaimids) that are primarily responsible for seed dispersal in the Asian tropics [19,23,24,38]. In species-rich communities, there is greater within-lineage diversity in key frugivore groups. For instance, across the six sites, the number of hornbill species varied from 0 to 7, bulbuls from 0 to 13, fruit-eating pigeons from 1 to 7 species, and barbets from 0 to 5. Among bird families that play a critical role in seed dispersal, such as Bucerotids, Pycnonotids, Sturnids, Columbids, Irenids, and Megalaimids, representatives were found between four to six of the sampled sites. Given the phylogenetic influence on traits, as demonstrated for two sites in an earlier study [10] and elsewhere [39], multiple species within a lineage are likely to have similar traits. This likely resulted in us not detecting any relationship between overall frugivore species richness at a site and community-weighted mean of beak width.

The inference of functional redundancy depends on the specific traits examined [40]. While the lack of a clear pattern in visitation rates and community-weighted mean of beak width suggests functional redundancy, other dimensions point towards functional complementarity. Like the strong association between beak and seed size, as also found in this study, highlights the dominant role of morphological trait matching in structuring plant-frugivore communities, which indicates functional complementarity. Moreover, closely related sympatric species may differ in the qualitative services they provide. For example, the Wreathed Hornbill (*Rhyticeros undulatus*) dispersed seeds over distances four times greater than its larger relative, the Great Hornbill (*Buceros bicornis*), in an Eastern Himalayan site also sampled in this study [41]. This has implications for seed dispersal in a changing world as Wreathed and Great Hornbills can play complementary roles with Wreathed Hornbill dispersing seeds over much larger distances (across fragments and/or wider elevational gradients), while Great Hornbills dispersing seeds locally. The absence of such sympatric species in species-poor sites may impact the qualitative role of seed dispersal function. Additionally, frugivores in species-rich sites have been observed to have narrower dietary niches, and these communities were less nested and more modular (in prep.) (also see [42]), reflecting increasing dietary specialisation and fidelity to fewer plant species after factoring for morphological constraints. This foraging fidelity likely arises from interspecific competition, as dietary plasticity and shifts in frugivore diets were observed in the species-poor site with fewer competitors [10]. However, whether frugivore loss in species-rich sites would lead to the relaxation of dietary niches and contribute to functional redundancy remains an open question.

Functional redundancy and complementarity strongly influence ecological resilience. Species-poor communities, characterised by the absence of closely-related sympatric species and low overall frugivore diversity, are more vulnerable to species extinctions due to a lack of ‘insurance effects’ [43,44]. For example, drastic reductions in Helmeted Hornbill (*Rhinoplax vigil*) populations over 20 years are correlated with increased mean densities of the Great Hornbills [45], likely compensating functionally for the loss of Helmeted Hornbills, as both species share similar diets dominated by figs and have comparable body sizes. Such potential functional insurance is often absent in species-poor sites. On Narcondam Island, the point-endemic Narcondam Hornbill (*Rhyticeros narcondami*) is the sole species feeding on large-seeded plants. Large-seeded species might face significant disadvantages if this species were to be lost. In contrast, species-rich sites, with greater lineage diversity and overall diversity of frugivores, are likely to be more resilient to species loss than species-poor communities. Thus, the ecological resilience of species-poor communities, which may not appear functionally compromised, needs to be evaluated vis-a-vis species-rich communities.

Despite being on the mainland, the frugivore community in the Western Ghats had frugivore species pool similar to that of the Andaman Island site and nearly half the diversity of the species-rich Eastern Himalayan site. Dispersal events strongly influence bird communities [46]. While herpetofauna and invertebrates in the Western Ghats exhibit high diversity and endemism due to in-situ speciation, bird diversity is strongly shaped by dispersal events over evolutionary time scales [47,48]. The isolation of the Western Ghats from the Himalaya and the north-east Indian hills, caused by the presence of extensive dry habitats in peninsular India and the shrinking of wet forest habitats over geological time scales, likely created barriers to bird dispersal thereby influencing the relatively depauperate species pool of the Western Ghats. As a result, the Western Ghats, despite being a mainland site resembles an island with a relatively low diversity of frugivores, a pattern that has also been reported for butterflies [49].

## 5. CONCLUSIONS

This study, based on extensive primary field observational data collected from six sites across mainland tropical Asia, addresses a critical gap in understanding the relationship between biodiversity and seed dispersal, a process that has received comparatively less attention. Unlike a previous study along a disturbance gradient [2], we found no positive association between frugivore richness and visitation rates and functional composition of frugivores across a richness gradient shaped by biogeographic processes. This is likely driven by niche-based community assembly processes and density compensation. While evidence of functional redundancy exists, the ecological resilience of the species-poor communities is likely to be low, highlighting the need for further exploration. Given the dietary shifts in frugivores in the absence of competitors, determining the influence of frugivore loss on the ecological resilience of communities is challenging. By comprehensively sampling plant-frugivore interactions across four biodiversity hotspots in mainland Asia, this study also addresses the Eltonian Shortfall [50]. Although recent species-specific studies have linked quantitative and qualitative seed dispersal measures to plant recruitment outcomes [51], our community-level, multi-site study precluded such analysis. However, measures like visitation rates and beak width that we have examined are known to be strongly linked to seed dispersal function. Future studies should compare recruitment patterns of fleshy-fruited plants in species-poor versus species-rich sites to understand the ecological implications of biodiversity loss.

## ETHICS

This was an observational study and did not require ethical approval from a human subject or animal approval committee.

## AUTHORS’ CONTRIBUTION

Conceptualisation: RN, JJ, NP, SG, AG; Methodology: RN; Formal Analysis: RN, JJ; Investigation: RN, SC, RM, AJ, AG, VS, SG, NP, ND, RK; Data curation: RN, SC, AG, RM, AJ, VS; Writing-Original Draft: RN; Writing-Review & Editing: All authors, Project Administration: RN, RM, AJ, BCS, VC, GG; Funding acquisition: RN, AJ, BCS, VC

### ACKNOWLEDGEMENTS

We thank the Forest Departments of Arunachal Pradesh, Andaman and Nicobar Islands, Tamil Nadu, and the Thailand Department of National Park, Wildlife and Plant Conservation. We thank Anand Osuri, Divya Mudappa, T. R. Shankar Raman, Aparajita Datta, Japang Pansa, Hasan Ali, Late Josna Malik, Devathi Parashuram, Daphawan Khamcha, Sitthichai Jinamoy, Siriwan Nakkuntod, Sunate Karapan, Sukanya Chaisuriyanun, and Narongsak Pongdee for support and critical discussions. We thank our field assistants across the different field sites: Pakke: Khem Thapa, Late Tali Nabam, Late Kumar Thapa, Andaman: Michael Kujur, Sujith Bengra, Namdapha: Dhan Bahadur Limbu, Wanggao Wangsa, Laiphong Wangnow, Anamalai: Krishnakumar, Sathiyaraj, and Bala: Isamaea Ma, Iswan Jowae, Masauphee Hub. We thank Siddhant Mehendale, Prabhav Benara, and Mandar Tijare for assisting us in data collection in Anamalais.

## DATA AVAILABILITY STATEMENT

Data will be uploaded on DataDryad post submission of the manuscript.

## SUPPLEMENTARY MATERIAL

**Figure S1.**
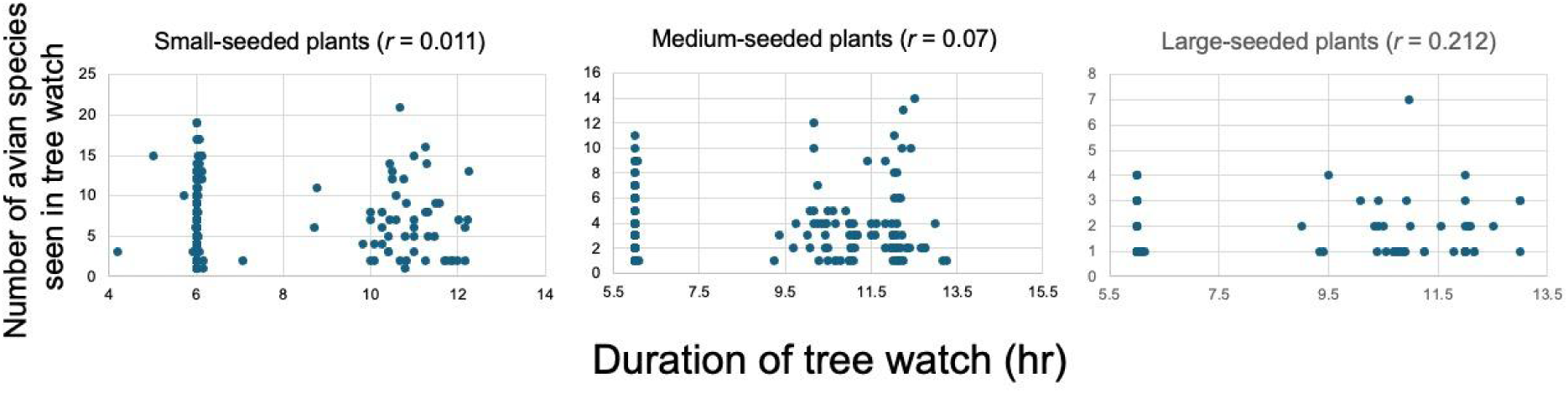
The duration of the tree watch did not influence the number of frugivore species detected on the fruiting trees across the three seed size class categories. Pearson’s correlation coefficient (*r*) is reported in parentheses.

**Figure S2.**
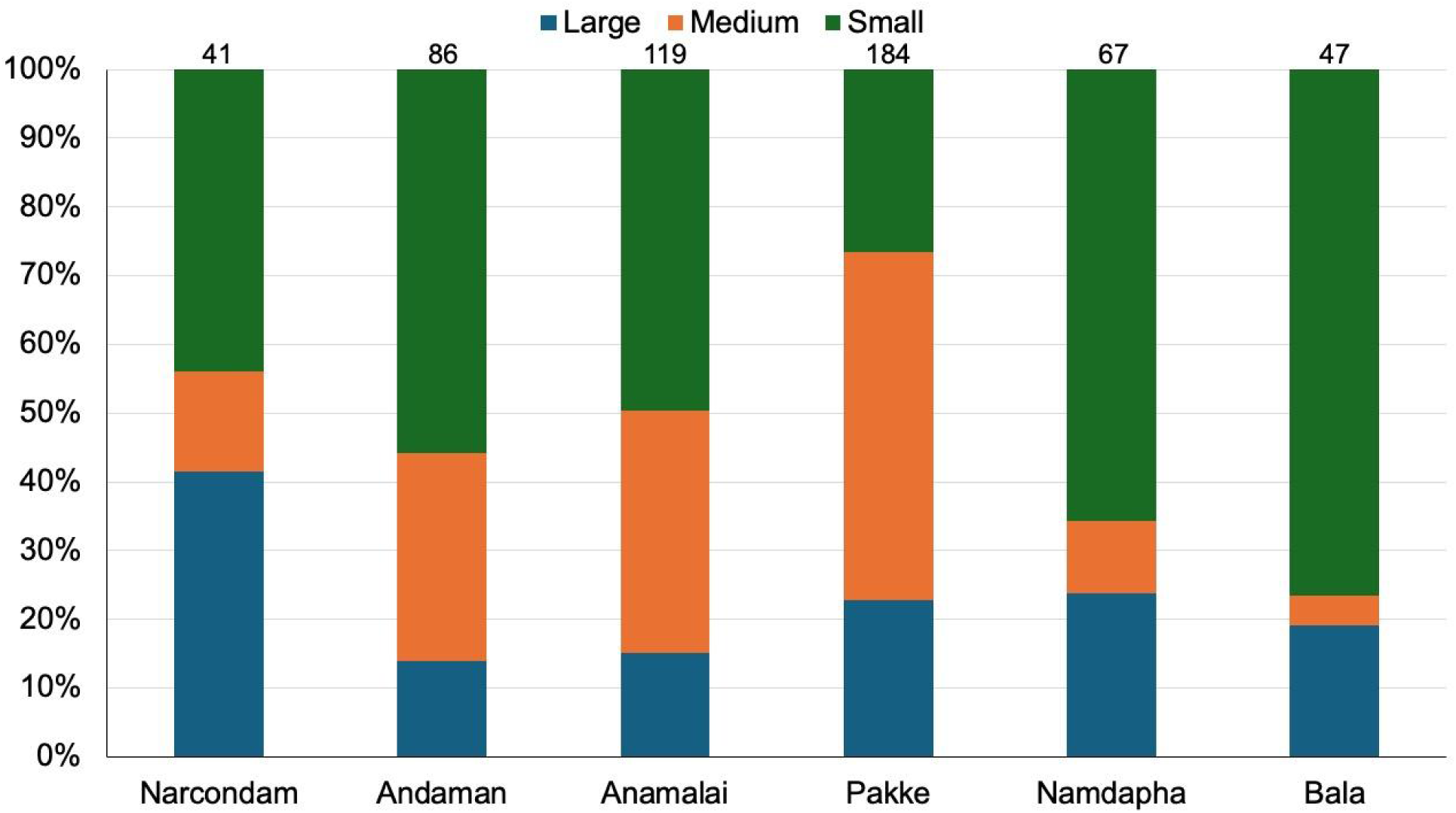
The proportion of trees with large, medium, and small-seeded plants observed in each site. The number in parentheses indicates the avian frugivore species detected at each site. The number on each bar represents the number of individual trees observed.

**Figure S3.**
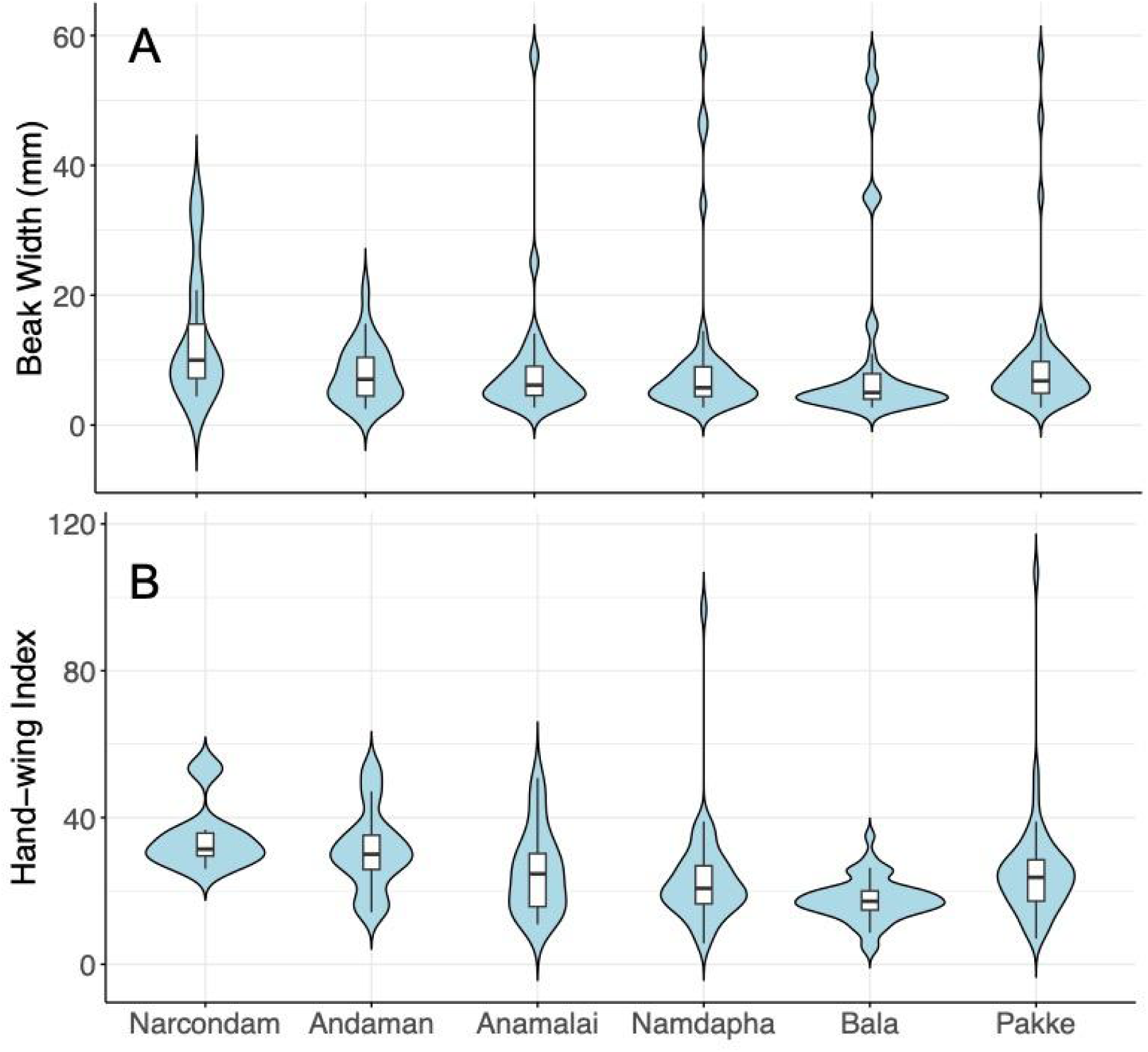
Violin plot showing the distribution of beak widths (A) and hand-wing index (B) across the six sites, arranged in increasing order of overall avian frugivore species richness in the community. The box plot nested within the violin plot depicts the median and the interquartile range.

**Table S1.**
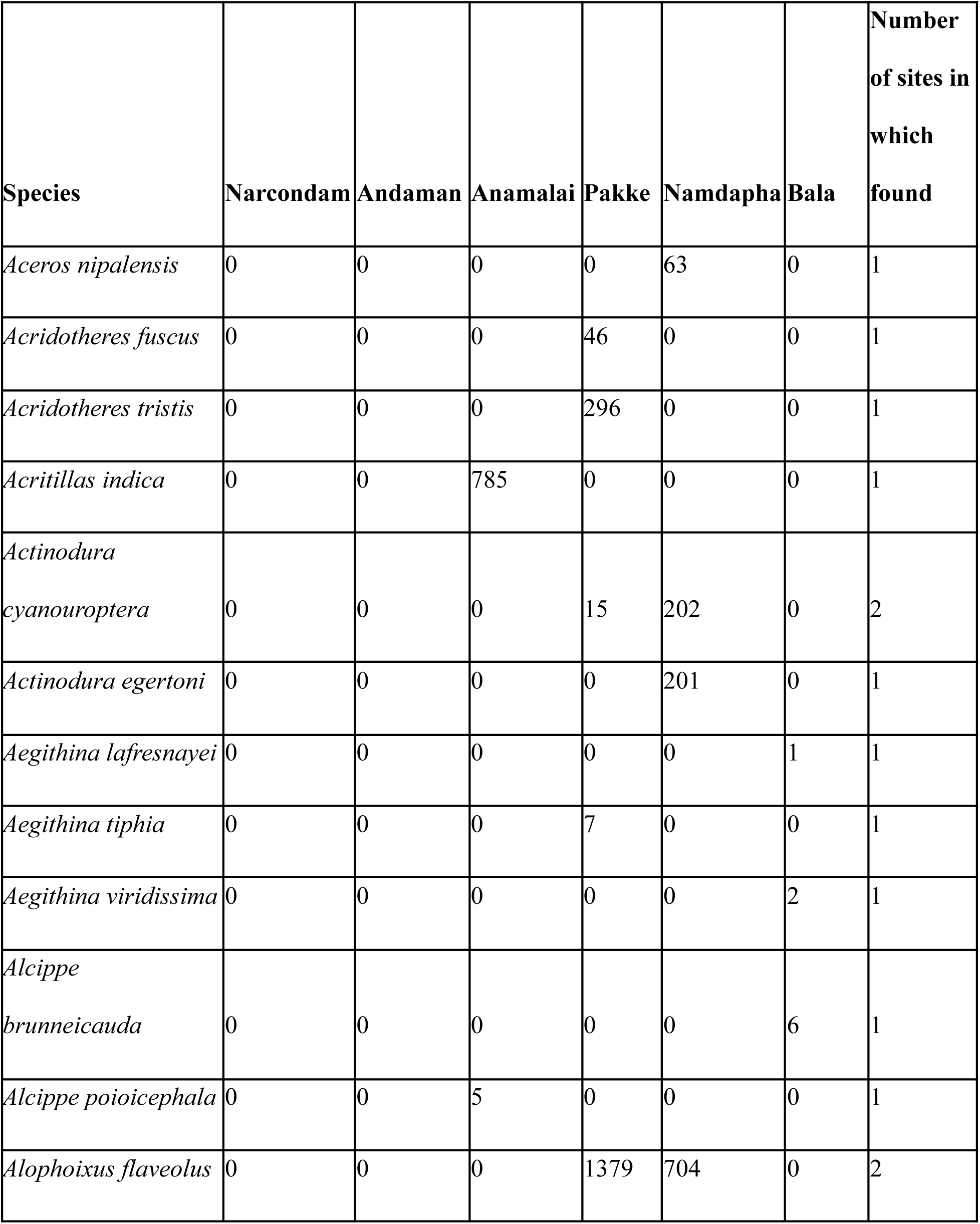
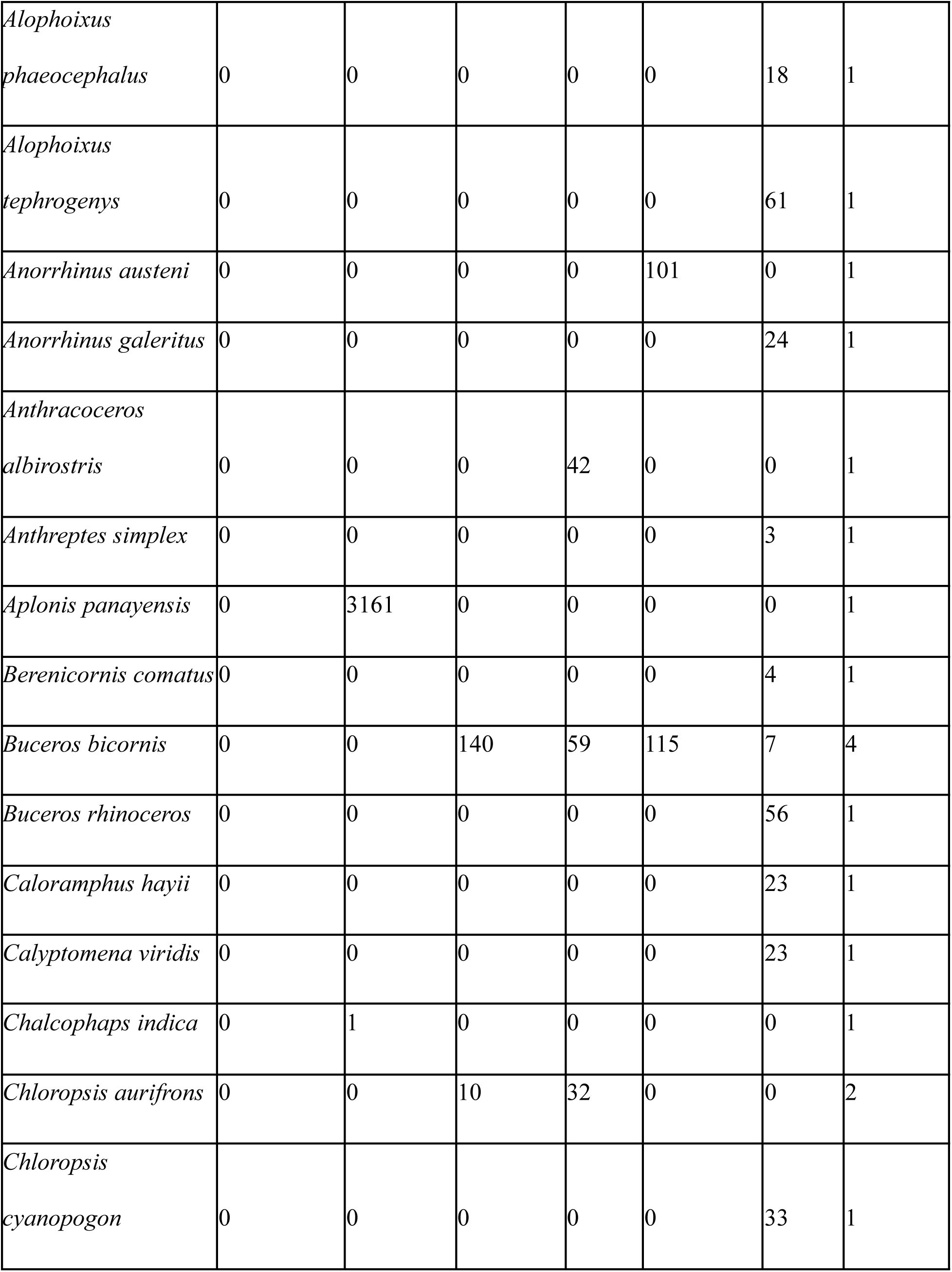
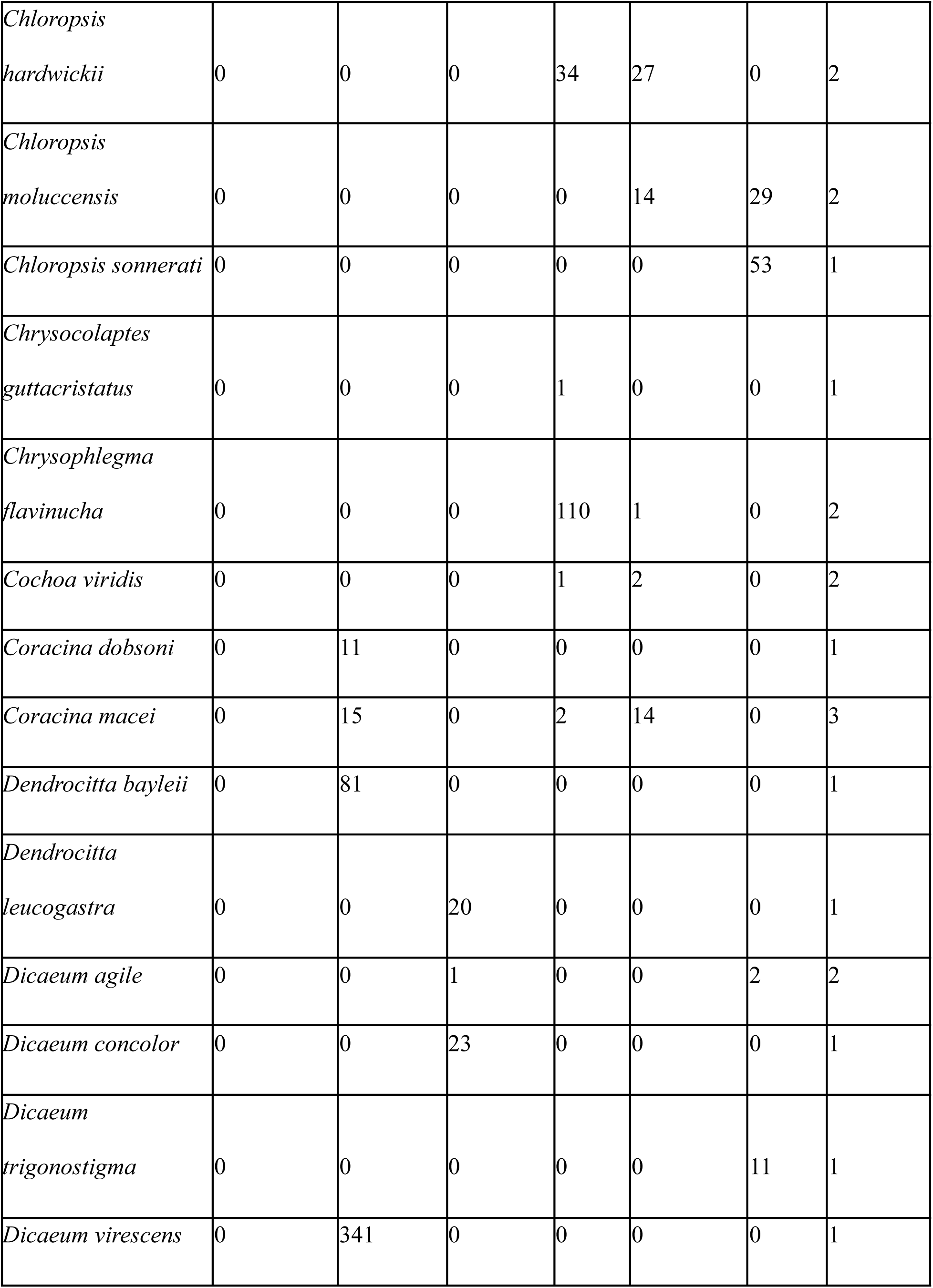
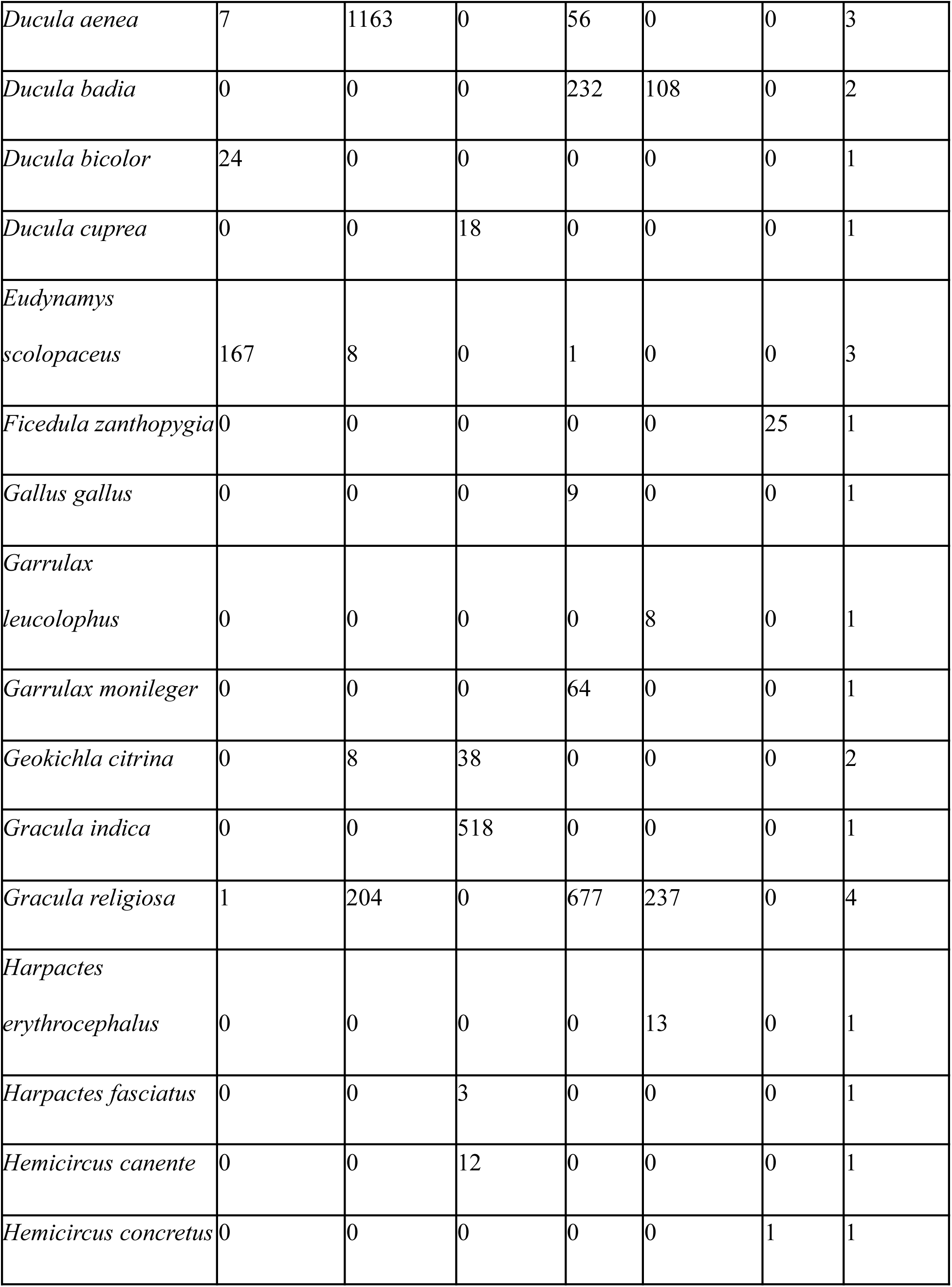
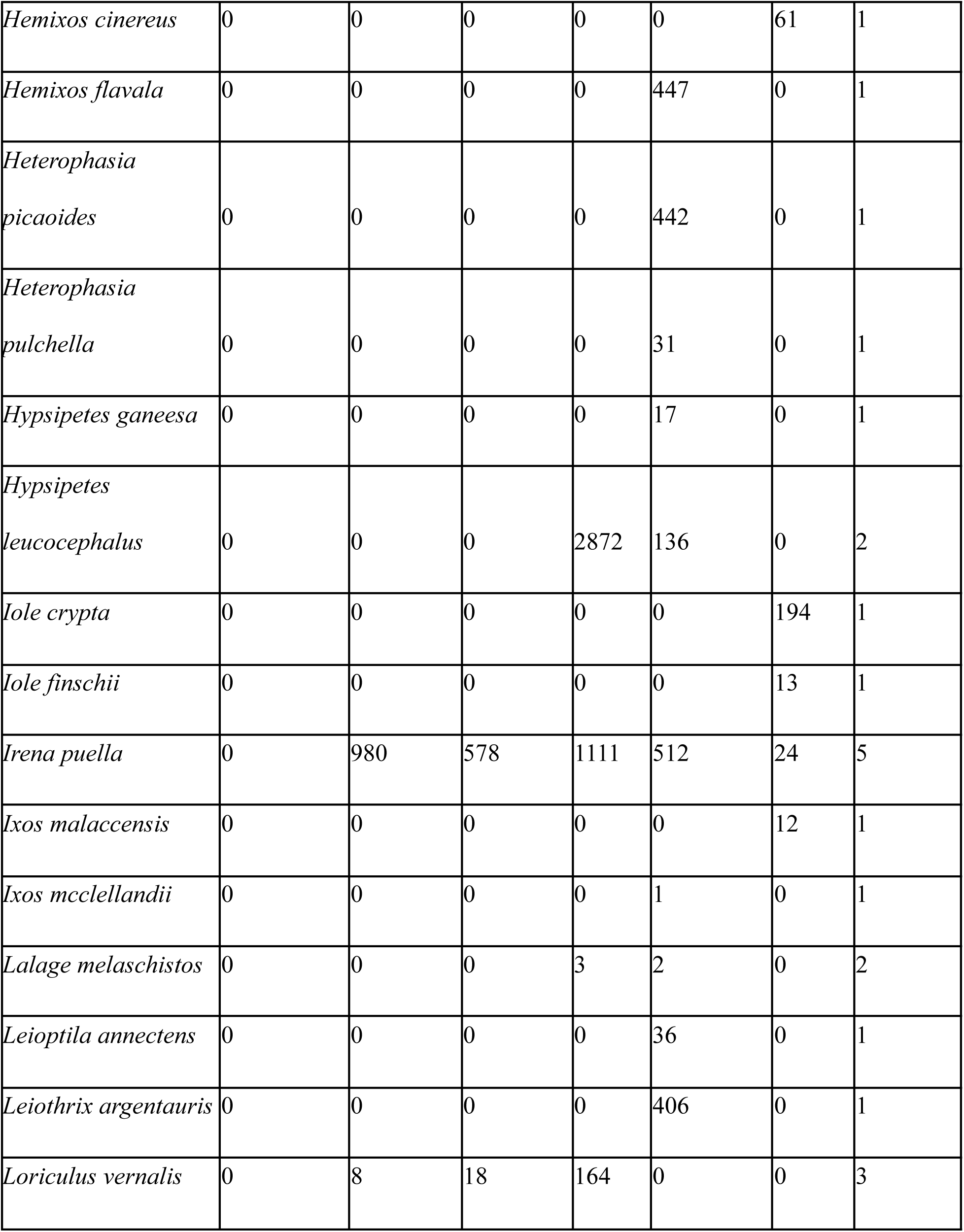
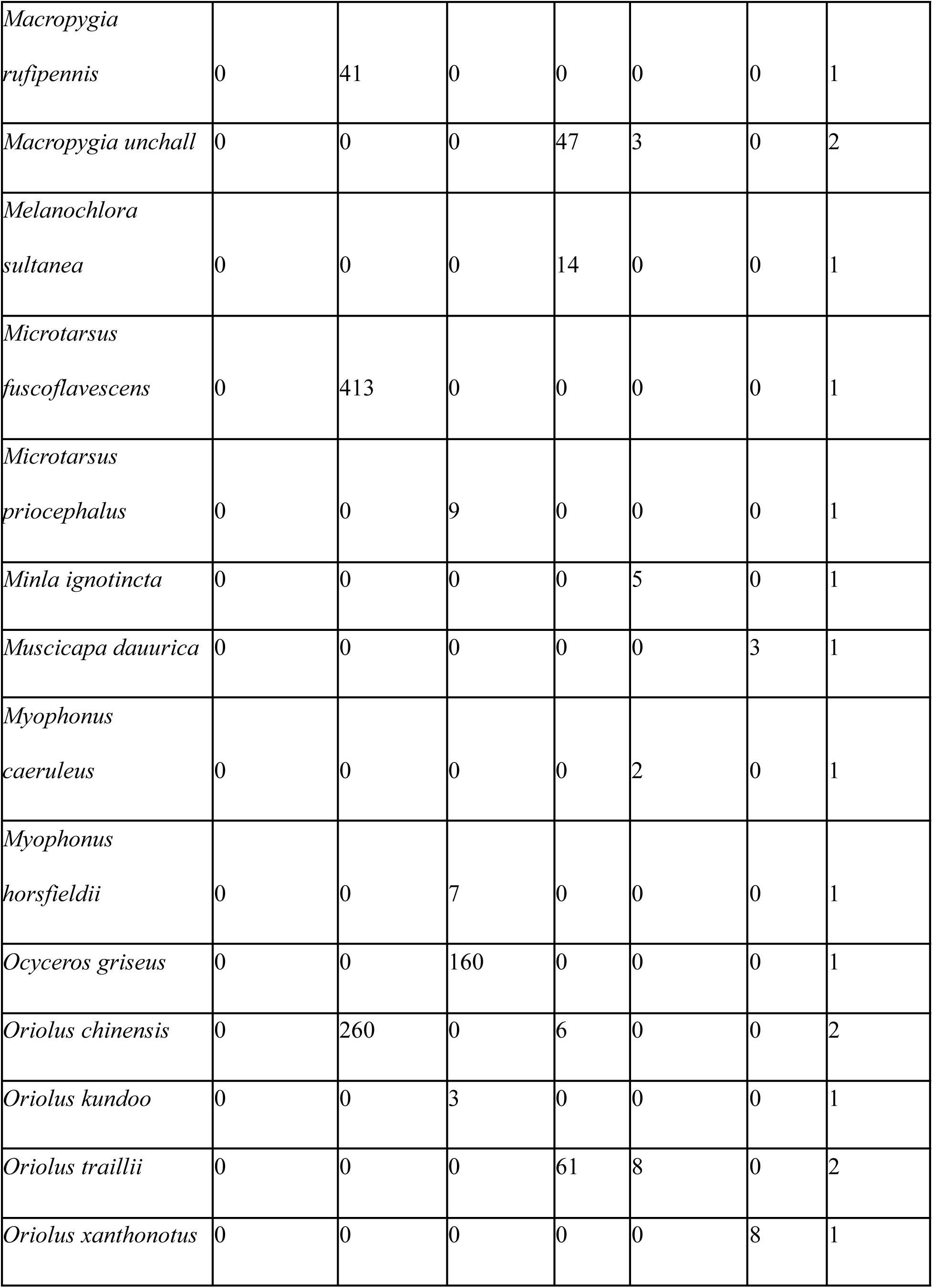
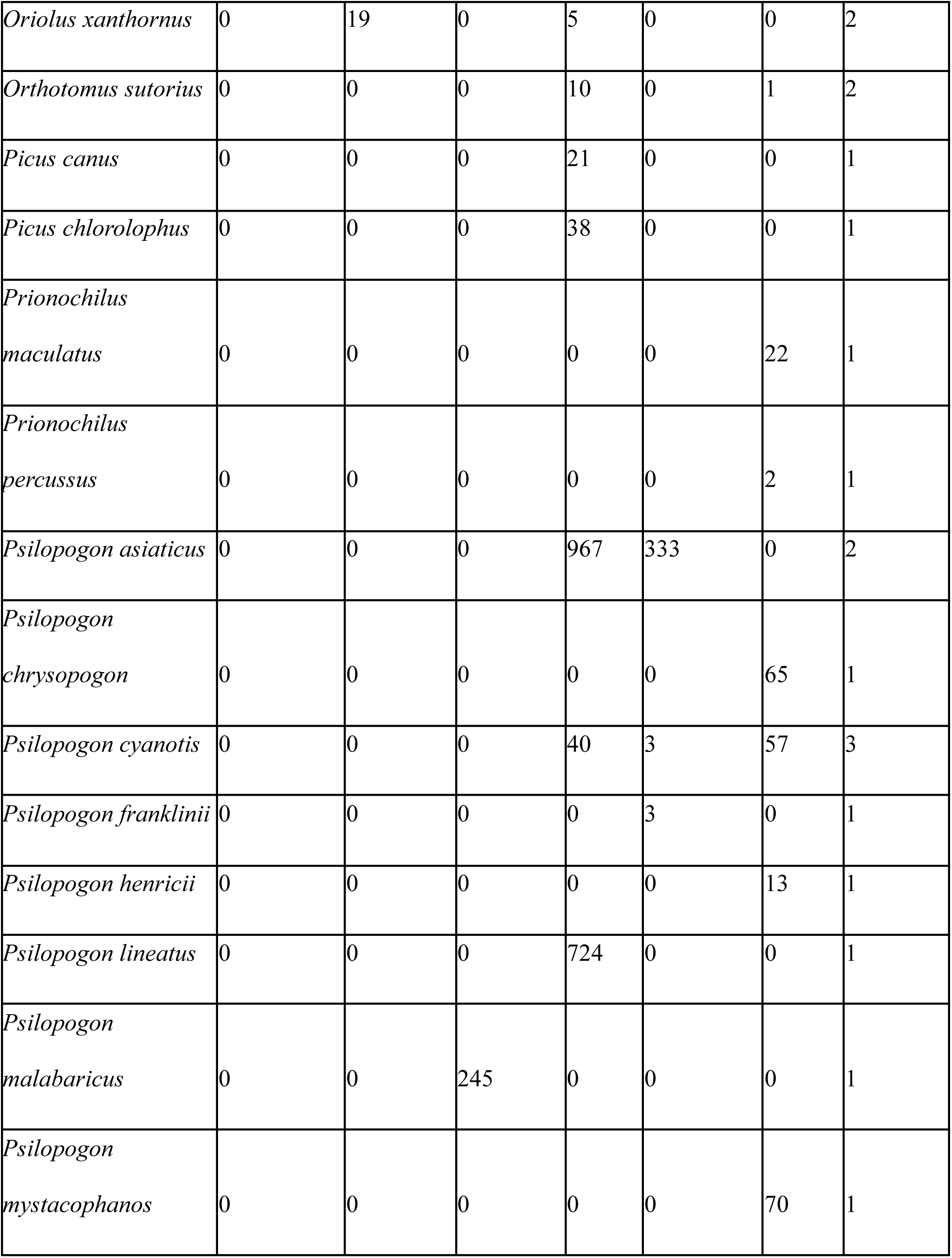
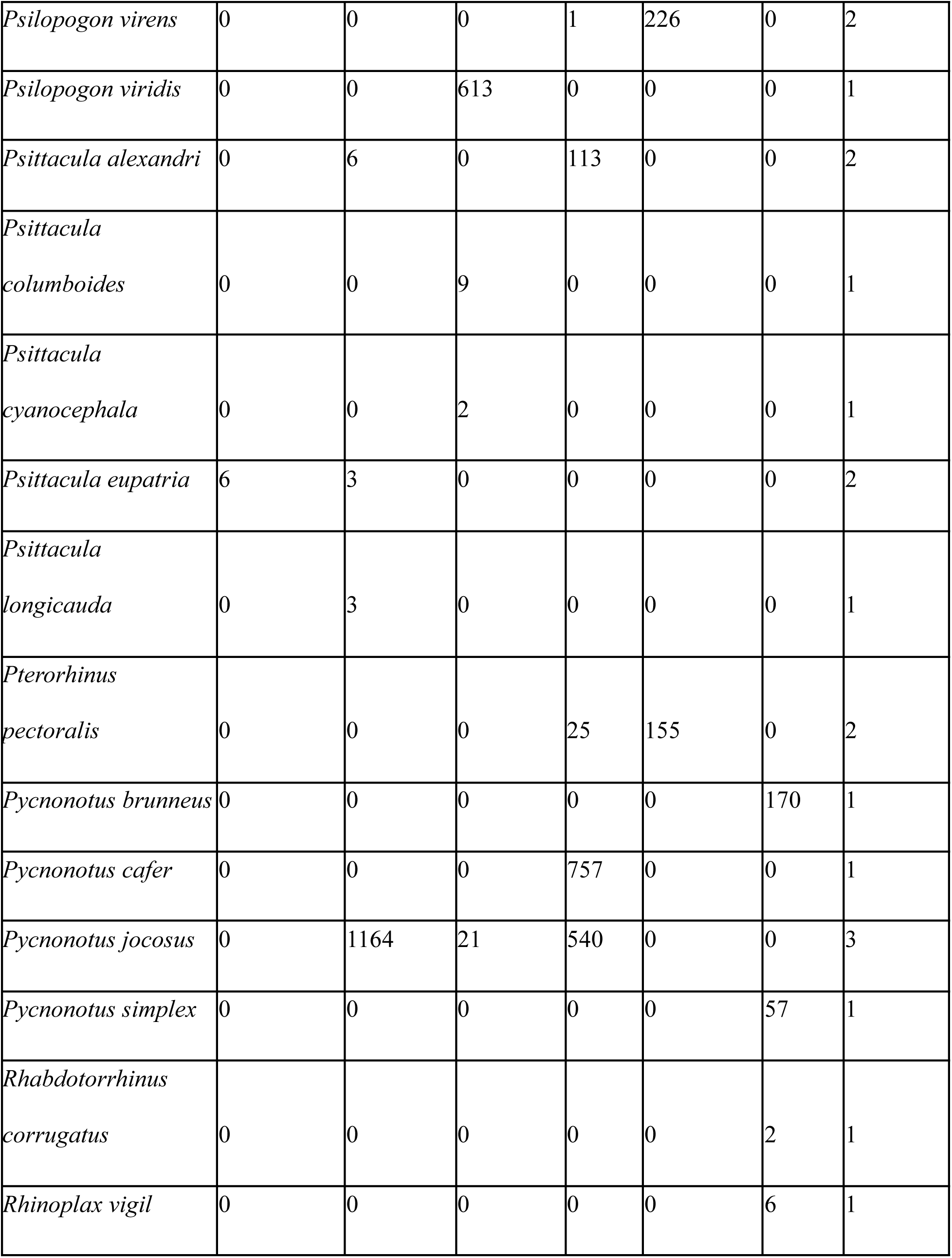
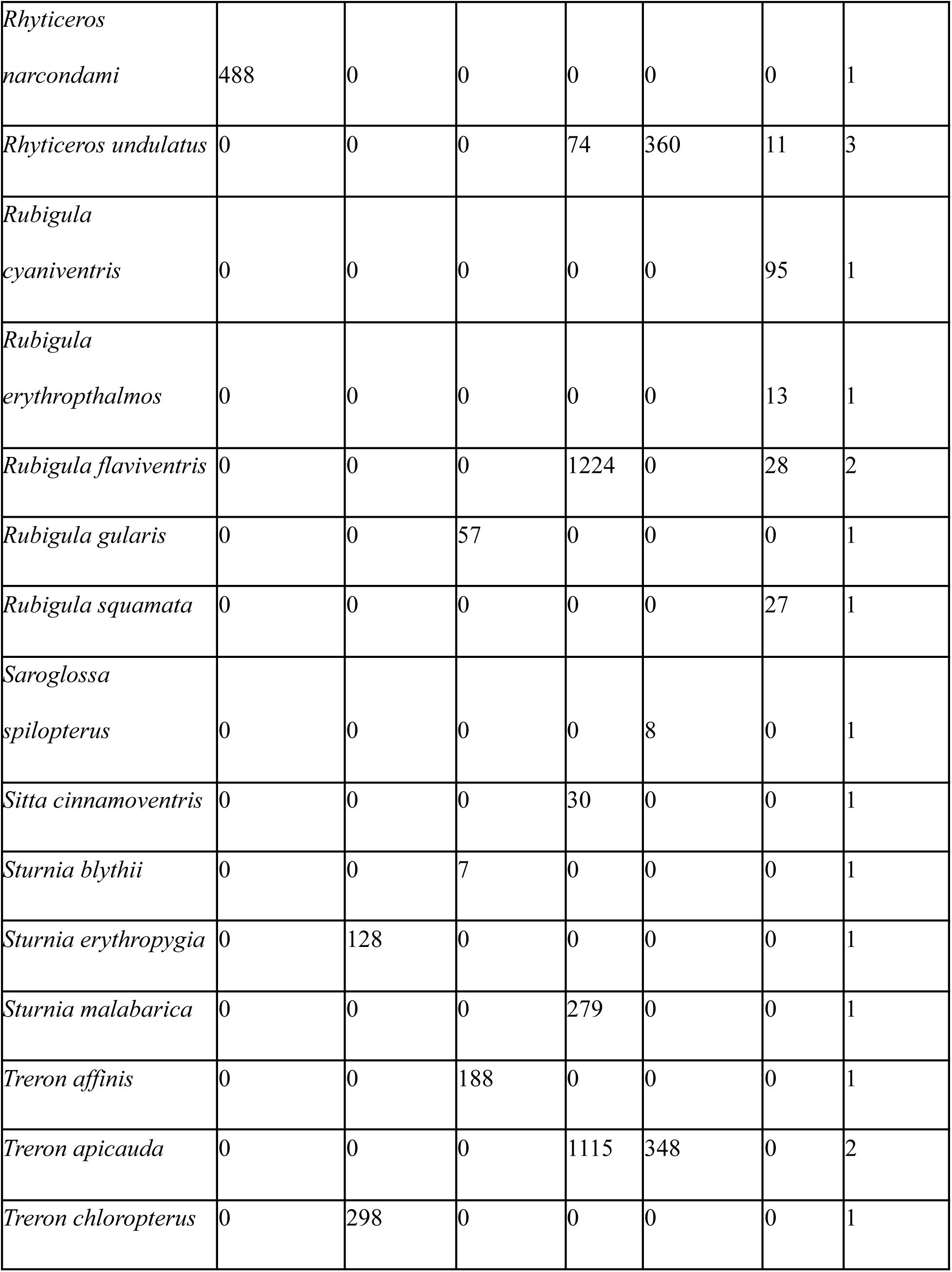
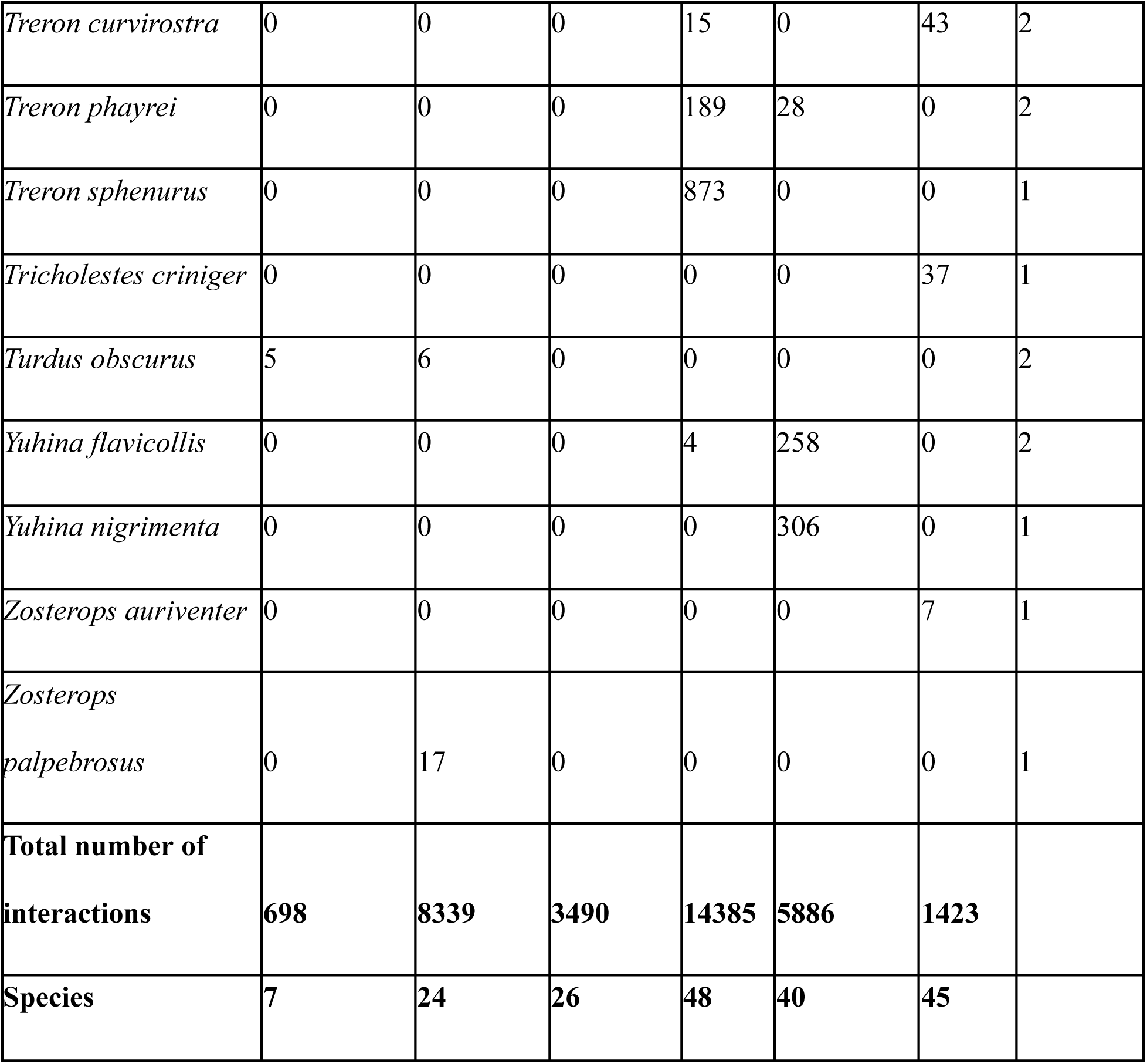
Number of individuals of avian frugivore species seen on fruiting trees in each of the six sites. The total number of interactions for each site and the number of unique avian frugivore species seen at each site is summarised at the bottom of the table. The number of sites where a particular species of frugivore was detected is summarised in the rightmost column.

**Table S2.** Details of sampling locations with the number of large-, medium- and small-seeded trees observed at each site.

|  |  |  | Large-seeded plants | Medium-seeded plants | Small-seeded plants | Total plant species observed |
| --- | --- | --- | --- | --- | --- | --- |
| Site name | Location | Biodiversity Hotspot |  |  |  |  |
| Narcondam | Andaman archipelago, India | Indo-Myanmar | 17 | 6 | 18 | 13 |
| Andaman | Andaman archipelago, India | Indo-Myanmar | 12 | 26 | 48 | 24 |
| Anamalai | Tamil Nadu, India | Western Ghats-Sri Lanka | 18 | 42 | 59 | 28 |
| Pakke | Arunachal Pradesh, India | Eastern Himalaya | 42 | 93 | 49 | 46 |
| Namdapha | Arunachal Pradesh, India | Indo-Myanmar | 16 | 7 | 44 | 28 |
| Bala | Narathiwat, Thailand | Sundaland | 9 | 2 | 36 | 22 |

**Table S3.**
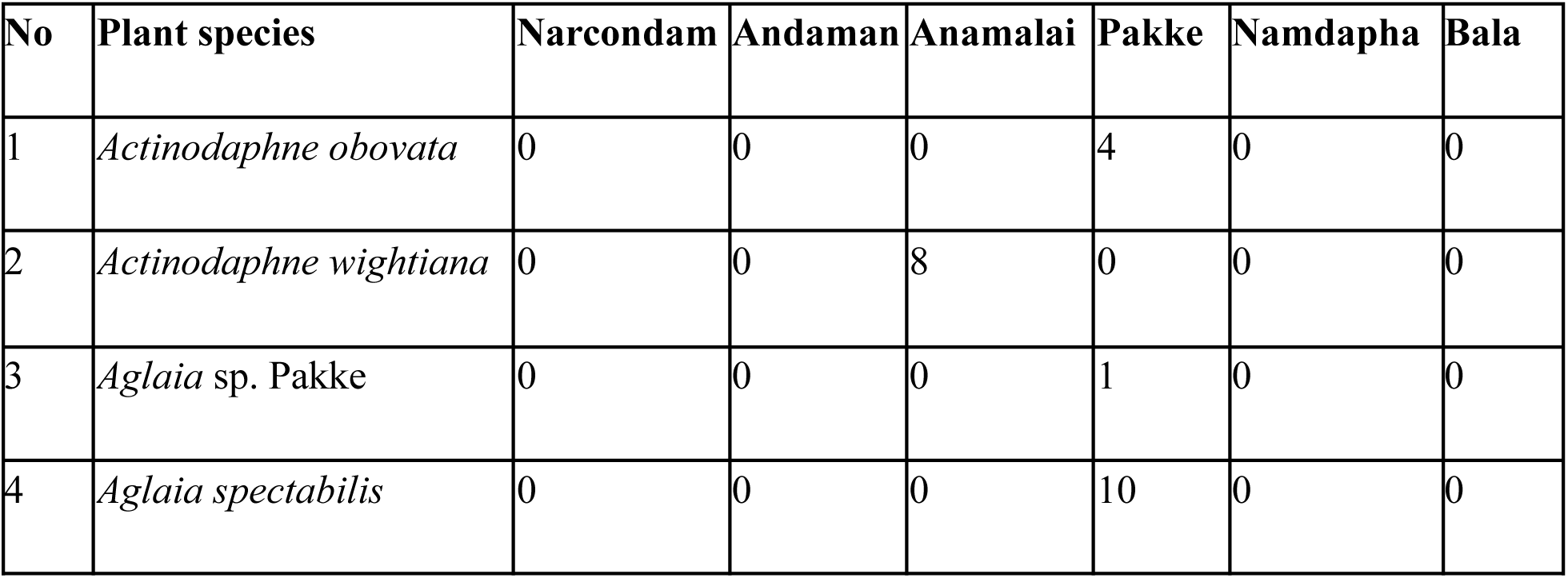
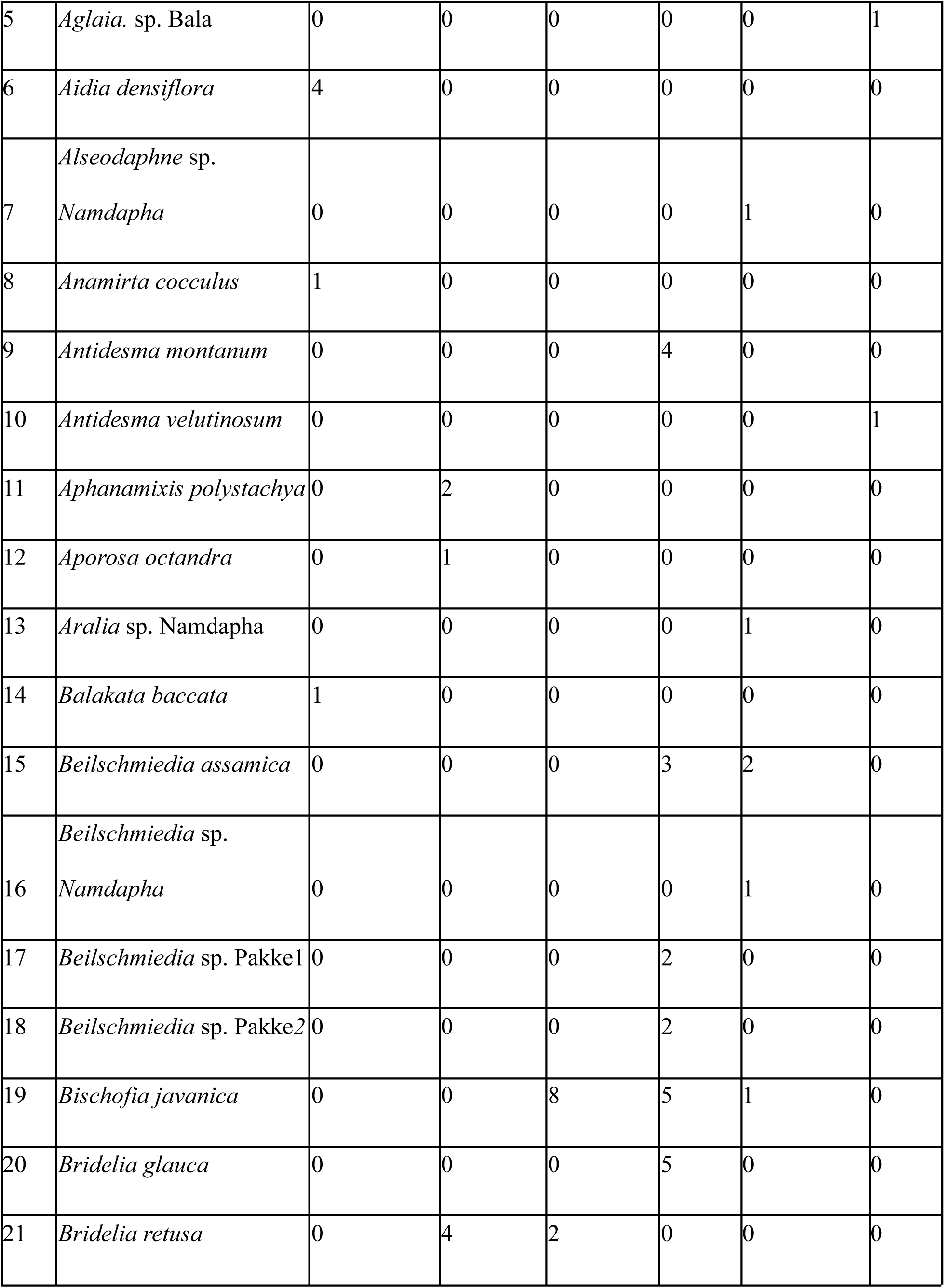
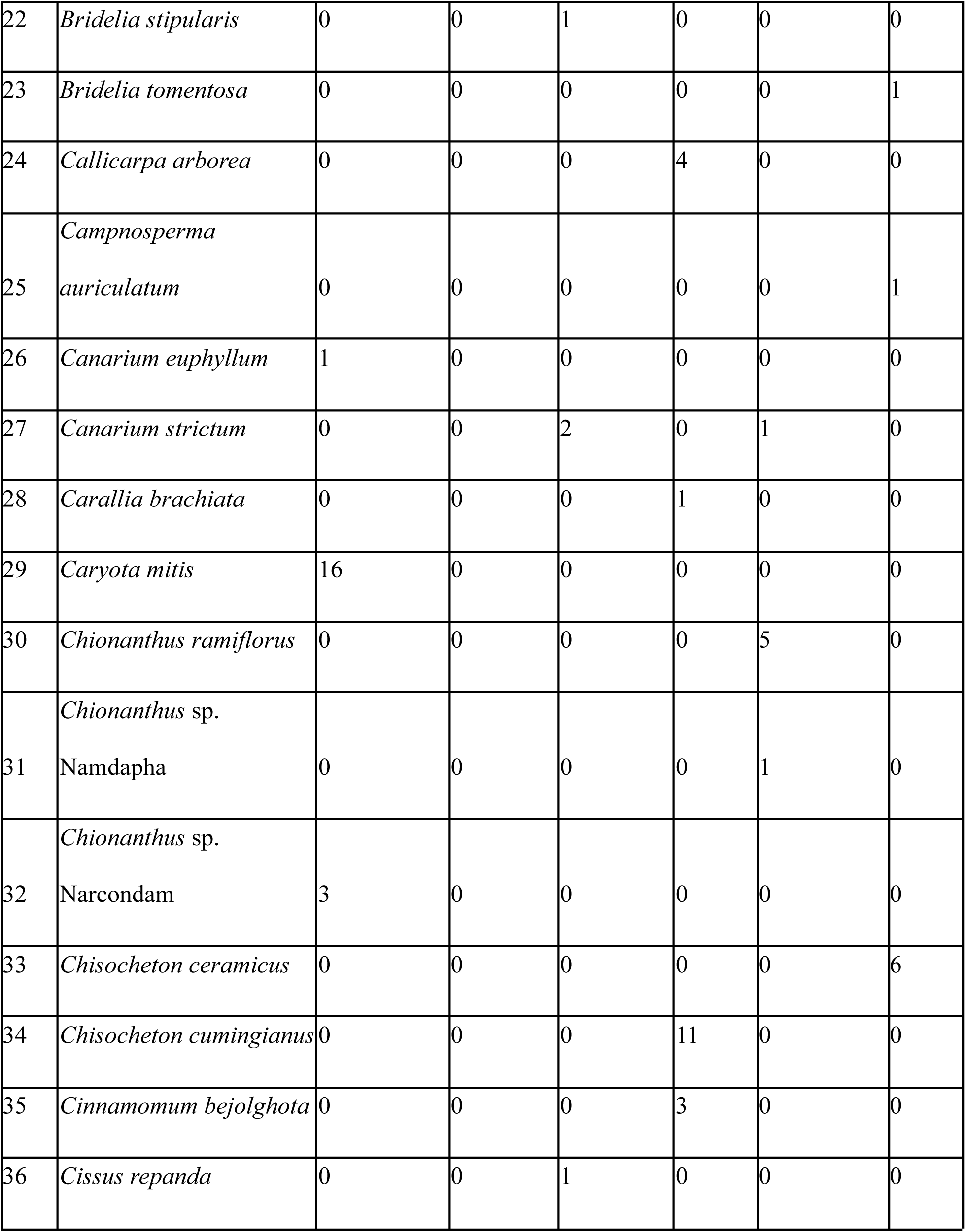
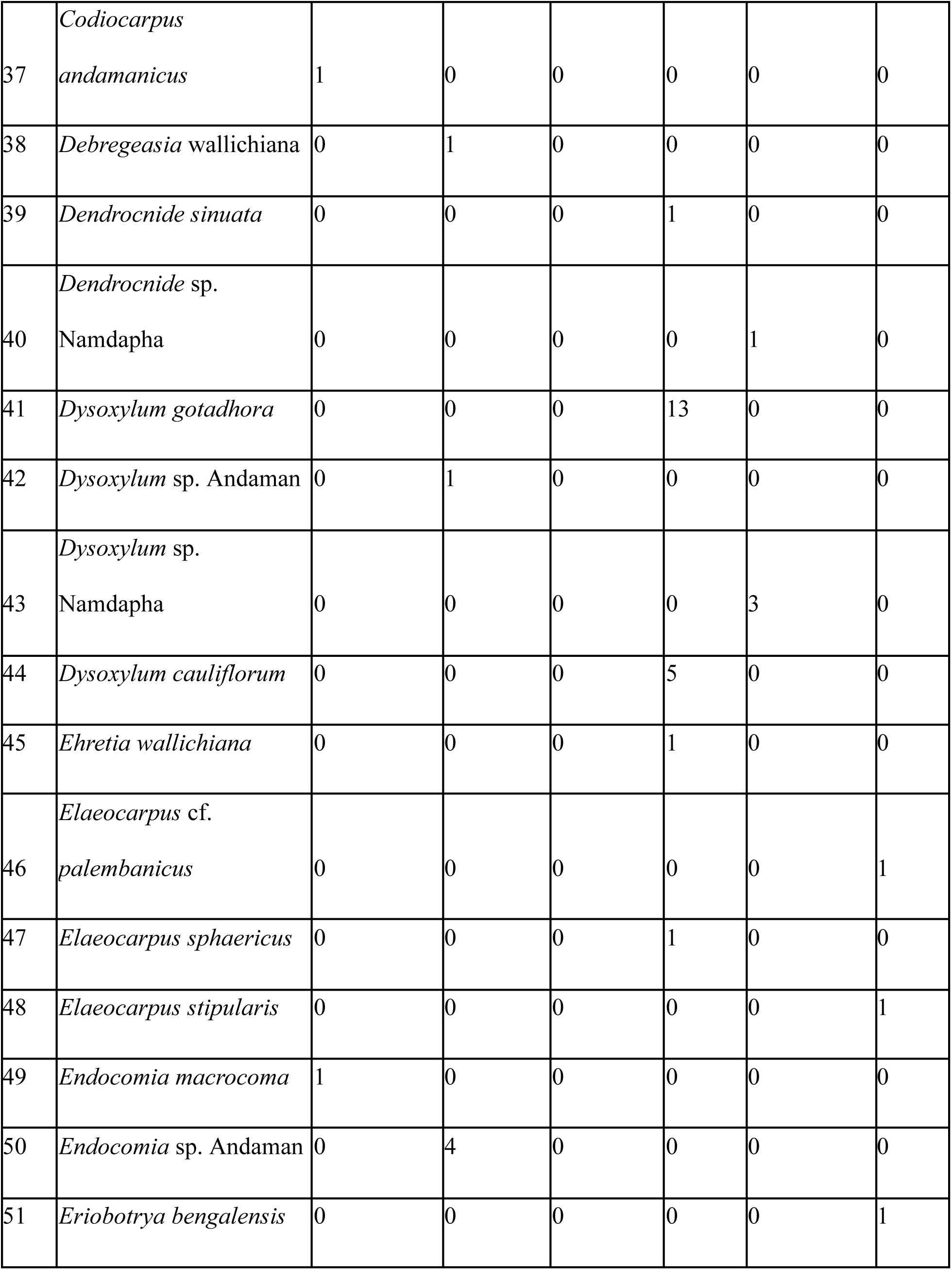
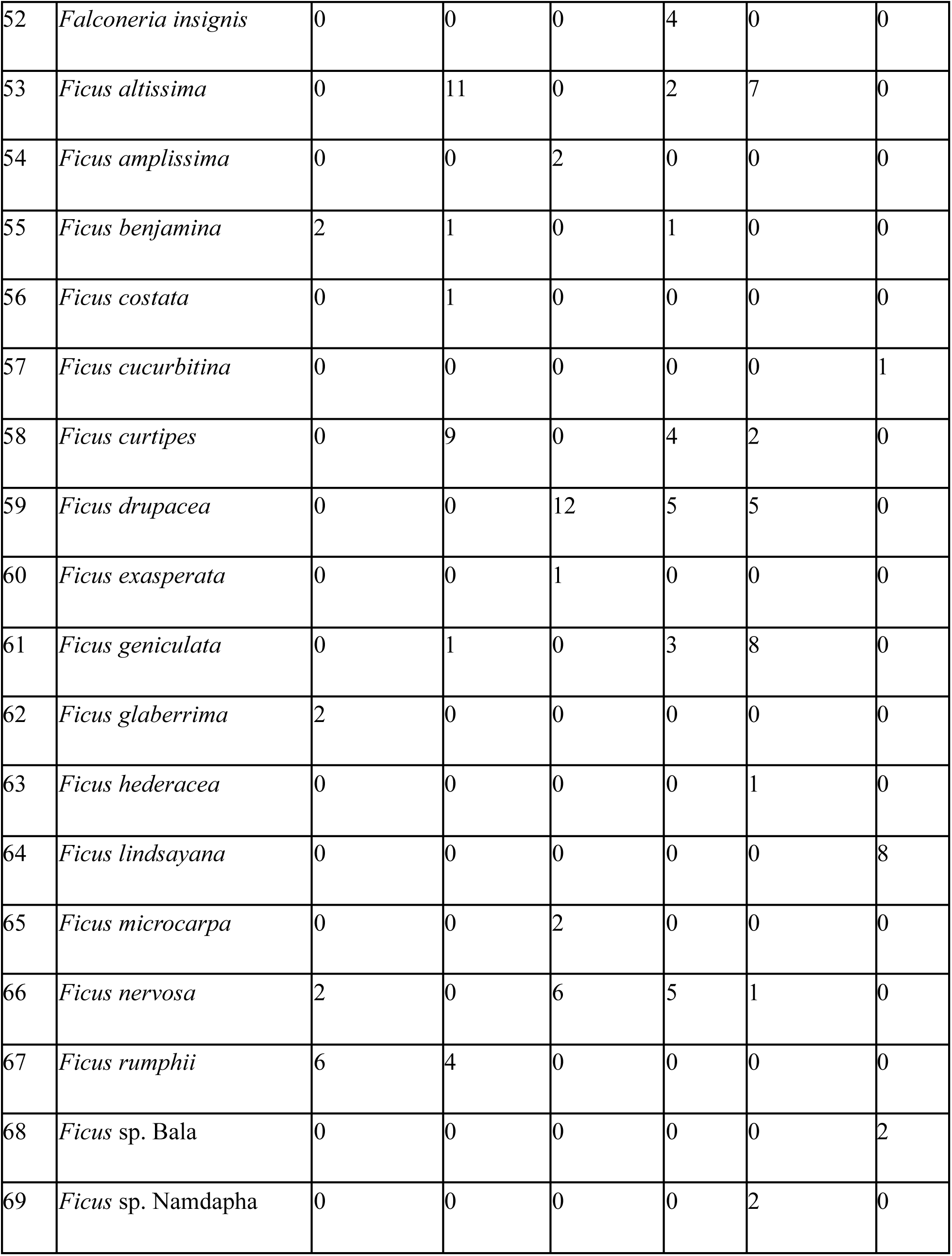
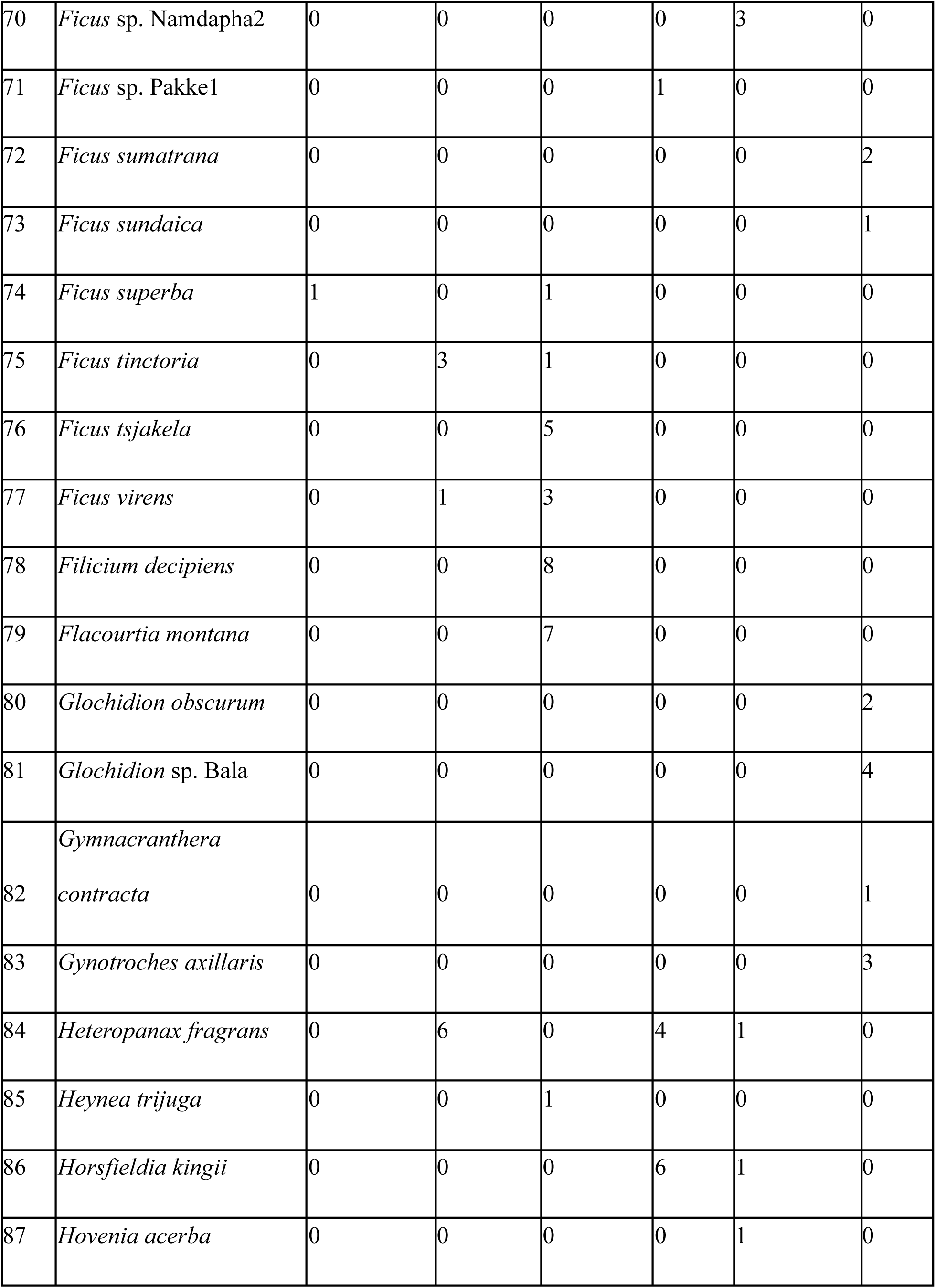
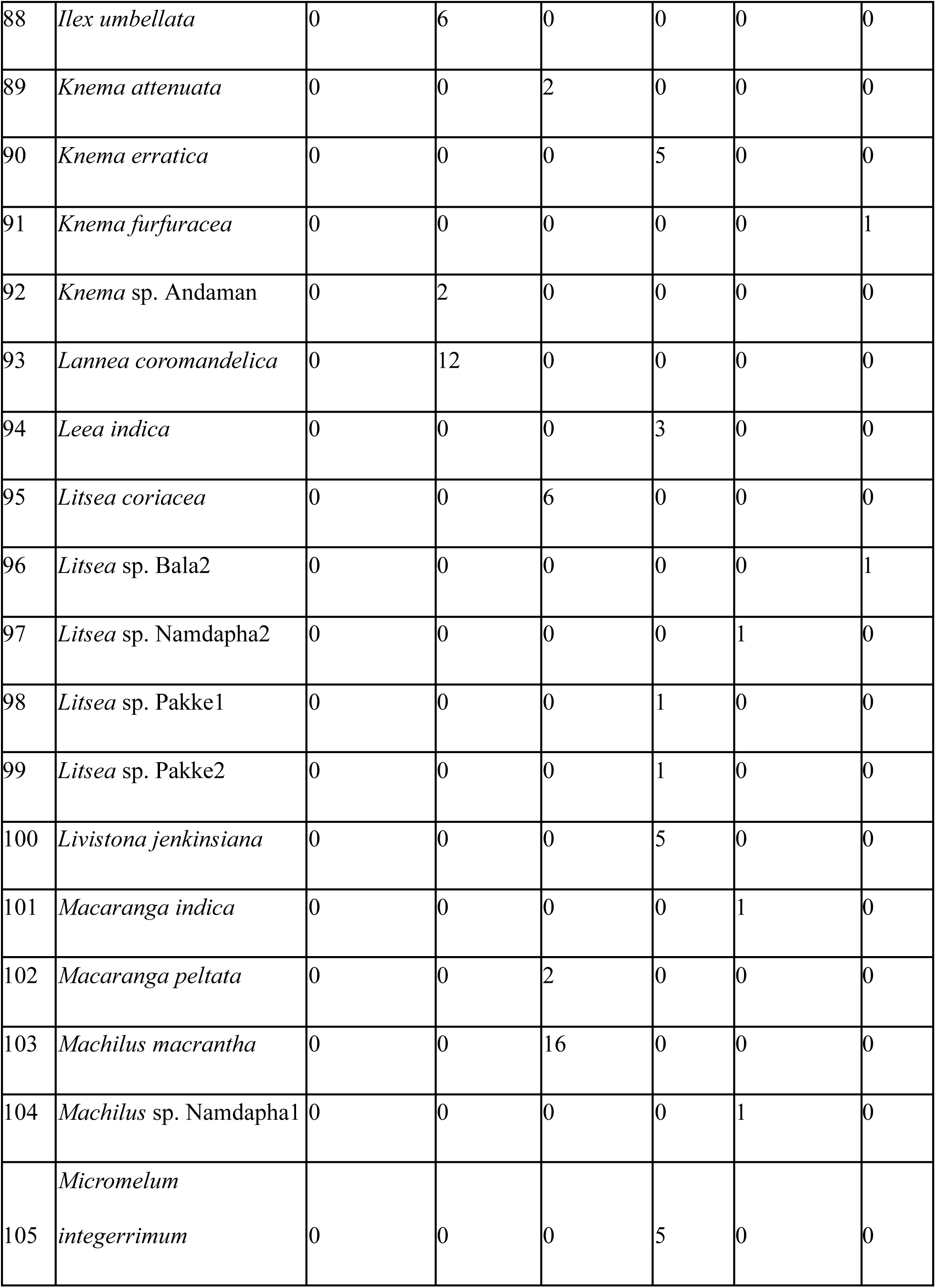
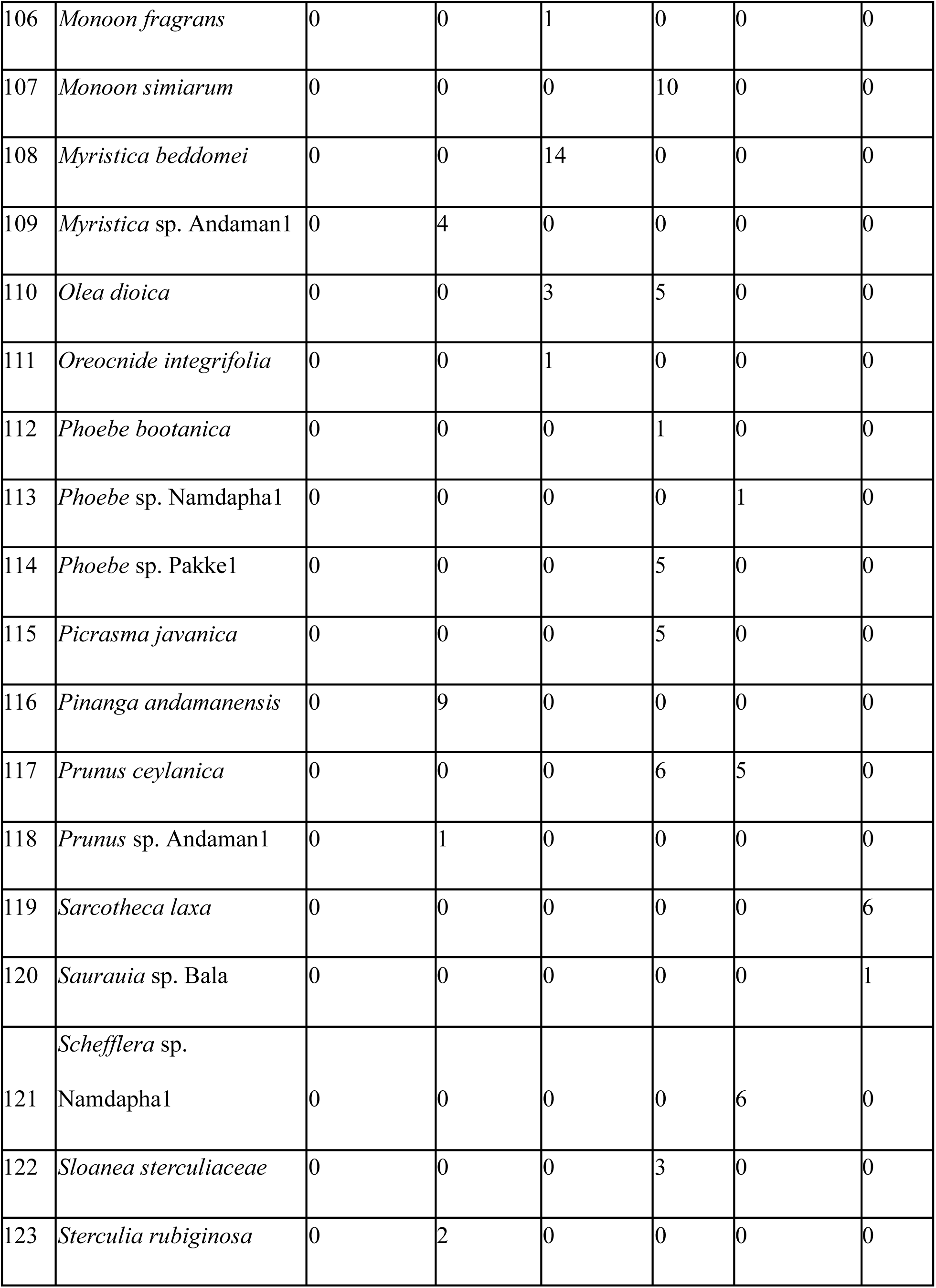
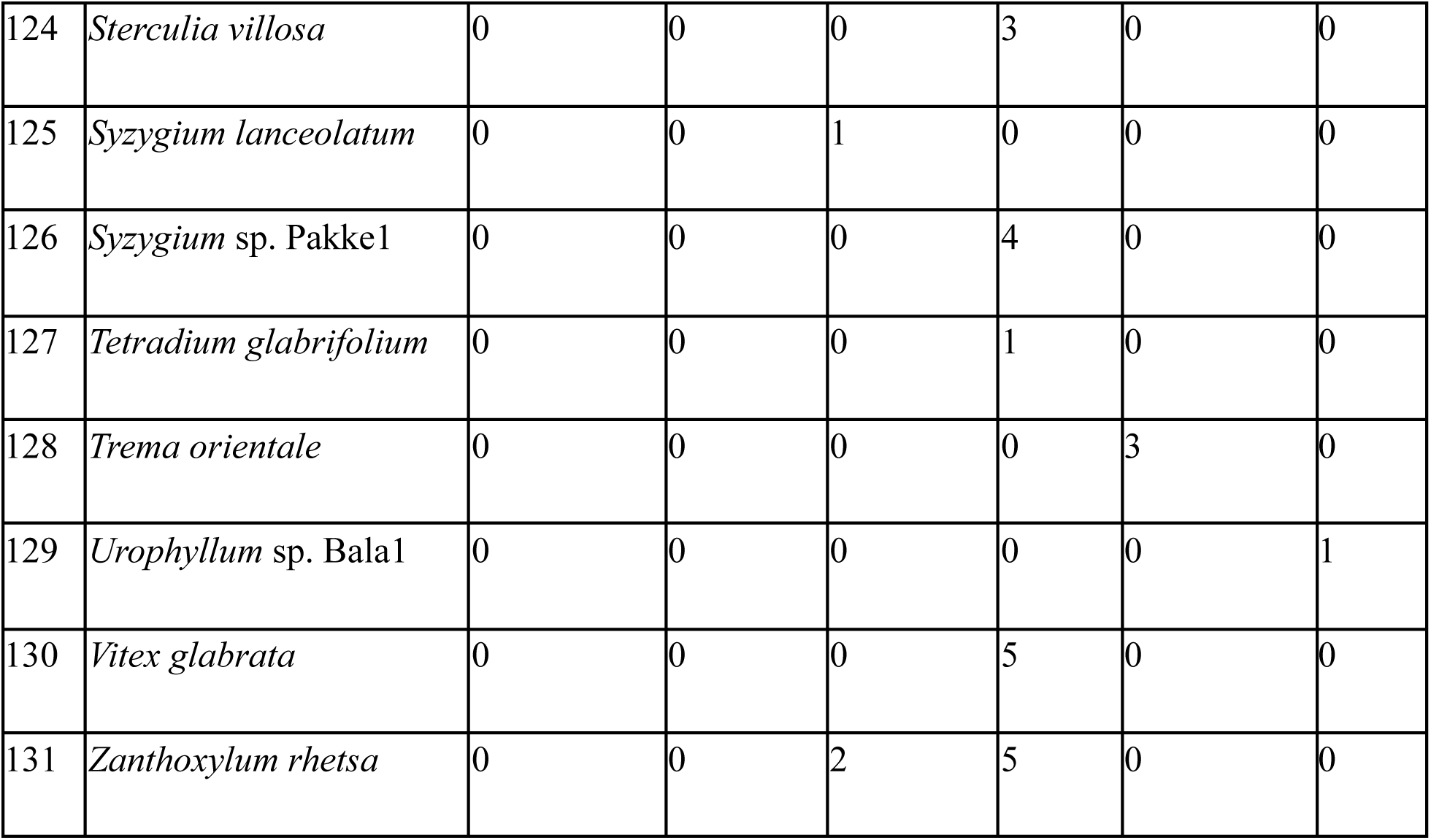
Number of individuals of different plant species that were observed across the six sites.

**Table S4.** Estimates of sampling coverage, overall species richness (qD), associated 95% CI and the overall avian frugivore species richness for each site.

| Site | Seed size | Sampling coverage | qD | 95% LCI | 95% UCI | Overall frugivore species richness |
| --- | --- | --- | --- | --- | --- | --- |
| Bala | Large | 0.944 | 4.8 | 3.3 | 6.4 | 45 |
| Pakke | Large | 1.000 | 11.0 | 11.0 | 11.0 | 48 |
| Narcondam | Large | 1.000 | 1.0 | 1.0 | 1.0 | 7 |
| Anamalai | Large | 1.000 | 3.0 | 2.6 | 3.4 | 26 |
| Namdapha | Large | 0.998 | 8.0 | 7.0 | 9.0 | 40 |
| Andaman | Large | 0.993 | 2.8 | 1.9 | 3.7 | 24 |
| Bala | Medium | 0.995 | 10.8 | 6.5 | 15.2 | 45 |
| Pakke | Medium | 1.000 | 45.6 | 33.7 | 57.5 | 48 |
| Narcondam | Medium | 0.997 | 4.4 | 1.3 | 7.5 | 7 |
| Anamalai | Medium | 1.000 | 18.4 | 14.4 | 22.4 | 26 |
| Namdapha | Medium | 1.000 | 13.2 | 9.6 | 16.9 | 40 |
| Andaman | Medium | 1.000 | 17.2 | 13.2 | 21.3 | 24 |
| Bala | Small | 0.995 | 42.1 | 38.7 | 45.5 | 45 |
| Pakke | Small | 1.000 | 43.1 | 30.4 | 55.8 | 48 |
| Narcondam | Small | 0.997 | 5.8 | 5.7 | 6.0 | 7 |
| Anamalai | Small | 1.000 | 24.3 | 20.2 | 28.4 | 26 |
| Namdapha | Small | 1.000 | 42.5 | 32.6 | 52.4 | 40 |
| Andaman | Small | 1.000 | 21.0 | 20.1 | 21.9 | 24 |

**Table S5.** Model summary table of the generalised linear mixed model (negative binomial error structure with site identity as the random effect) that examined the influence of overall avian frugivore species richness in the community and seed size (small, medium and large) on the number of avian frugivore species richness that visited a fruiting tree.

|  | Estimate | SE | z | p |
| --- | --- | --- | --- | --- |
| **Intercept (Seed size - Large)** | **0.456** | **0.113** | **4.046** | **< 0.001** |
| **Seed size - Medium** | **0.817** | **0.100** | **8.138** | **< 0.001** |
| **Seed size - Small** | **1.416** | **0.094** | **15.103** | **< 0.001** |
| **Overall species richness (centered)** | **0.023** | **0.006** | **3.883** | **< 0.001** |
| Random effects | Variance | SD |  |  |
| Intercept (Site) | 0.030 | 0.172 |  |  |

**Table S6.**
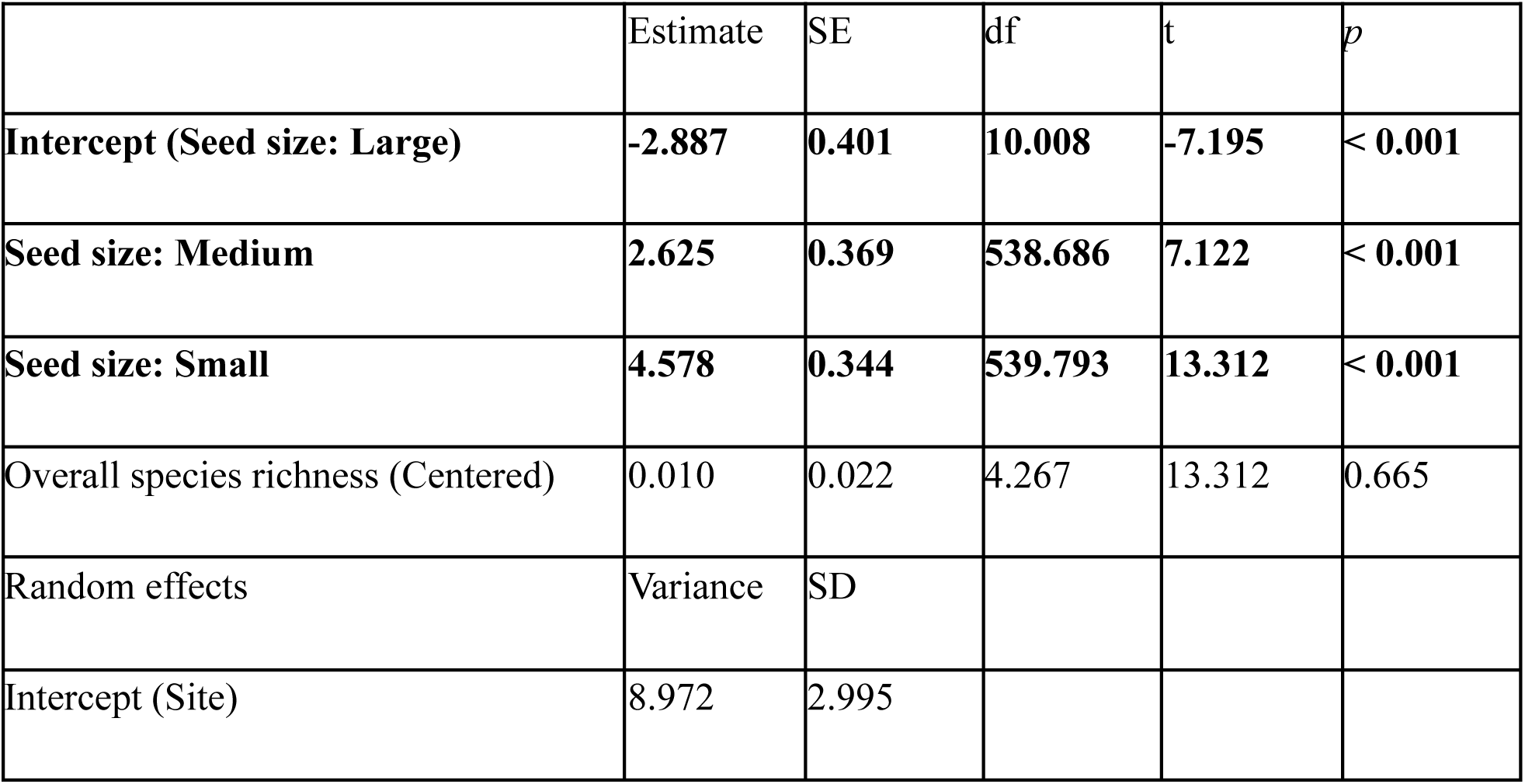
Model summary table of the general linear model (Gaussian error structure) that examined the influence of overall avian frugivore species richness in the community and seed size (small, medium and large) on the natural logarithm of visitation rates of frugivores that visited a fruiting tree during the tree watch. Overall species richness was centered to reduce strong correlation between that and the intercept. The significant parameters have been highlighted in bold.

**Table S7.** Model summary table of the general linear model (Gaussian error structure) that examined the influence of overall avian frugivore species richness in the community and seed size (small, medium and large) on the community weighted mean of the beak width (log-transformed for approximating normality) of all the frugivores that visited a fruiting tree during the tree watch. Overall species richness was centered to reduce strong correlation between that and the intercept. The significant parameters have been highlighted in bold.

|  | Estimate | SE | df | t | p |
| --- | --- | --- | --- | --- | --- |
| **Intercept (Seed size: Large)** | **1.055** | **0.069** | **5.083** | **15.229** | **< 0.001** |
| **Seed size: Medium** | **-0.297** | **0.033** | **479.426** | **-9.054** | **< 0.001** |
| **Seed size: Small** | **-0.391** | **0.030** | **479.039** | **-12.881** | **< 0.001** |
| Overall species richness (Centered) | -0.006 | 0.004 | 4.062 | -1.357 | 0.245 |
| Random effects | Variance | SD |  |  |  |
| Intercept (Site) | 0.023 | 0.153 |  |  |  |

**Table S8.**
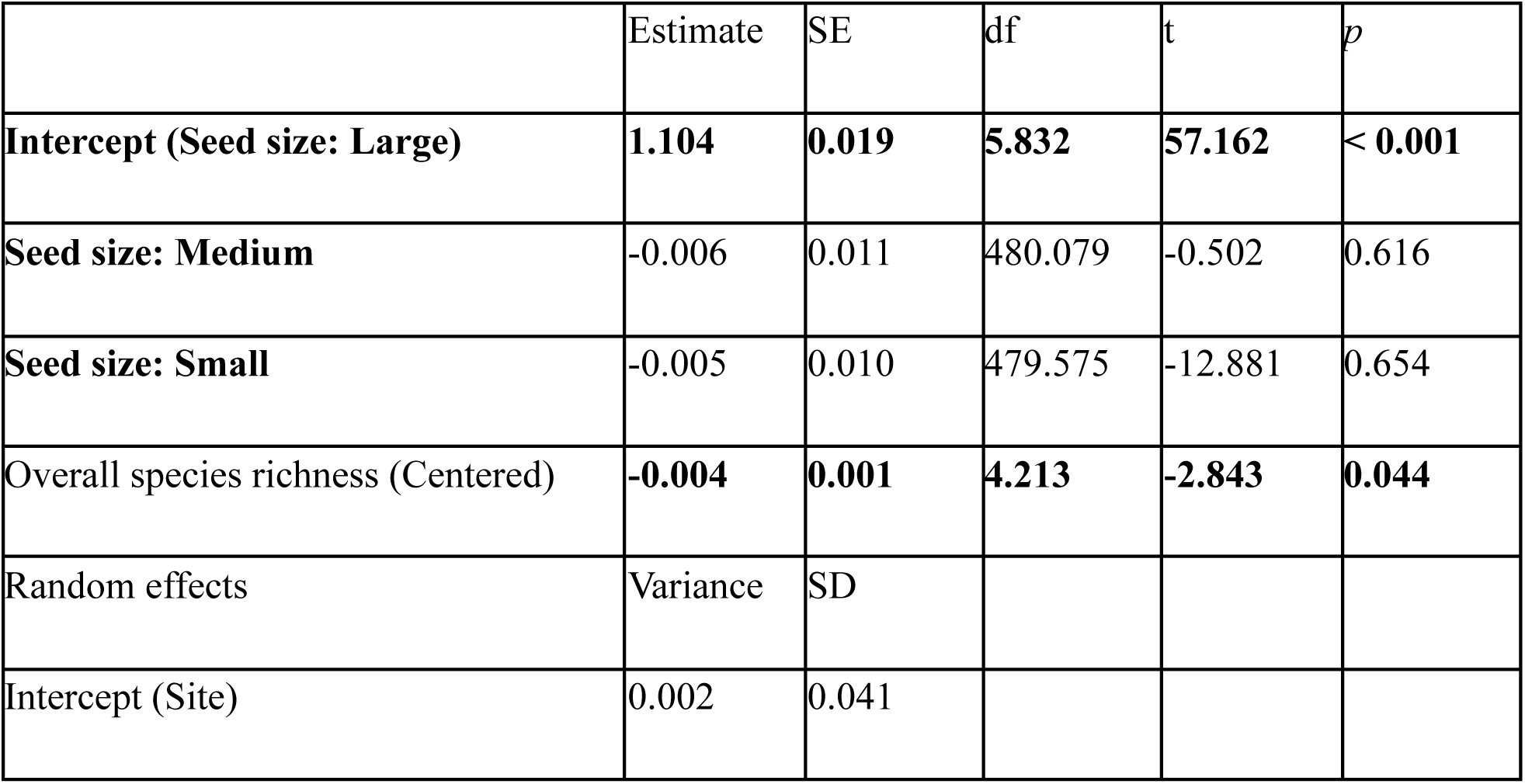
Model summary table of the general linear model (Gaussian error structure) that examined the influence of overall avian frugivore species richness in the community and seed size (small, medium and large) on the community weighted mean of the hand-wing index (log-transformed for approximating normality) of all the frugivore that visited a fruiting tree during the tree watch. Overall species richness was centred to reduce the strong correlation between that and the intercept. The significant parameters have been highlighted in Bold.

